# Modelling multicellular coordination by bridging cell-cell communication and intracellular regulation through multilayer networks

**DOI:** 10.64898/2026.01.20.700561

**Authors:** Rémi Trimbour, Ricardo O. Ramirez Flores, Julio Saez-Rodriguez, Laura Cantini

**Author notes:** These two authors jointly supervised this work (Julio Saez-Rodriguez, Laura Cantini).

## Abstract

In multicellular organisms, cells with various roles and locations coordinate to provide systemic and cohesive response to perturbations. These complex behaviors emerge from a complex interplay between intracellular regulation and intercellular signals that mediate cell-cell communication. While single-cell technologies opened the possibility of studying both, most methods focus solely on one of these aspects. Thus, they are only able to partially recover in vivo and multicellular behaviors.

We here introduce ReCoN (REconstruction of multicellular COordination Networks from single-cell data), a framework combining intracellular gene regulation and cell-cell communication to provide insights into multicellular coordination from single-cell data. First, ReCoN infers from single-cell data a heterogeneous multilayer network containing both cell-type-specific intracellular subnetworks and ligand-receptor interactions. Through random walk with restart explorations, ReCoN then infers the response of each cell type to both intra- and extracellular perturbations, such as a gene knock-out or a cytokine, respectively.

ReCoN was evaluated on predicting the *in vivo* response of immune cell-types to different cytokines and on recovering cardiac cell-type response in heart failure. It highlighted the role of indirect effects, where cells emit secondary messengers in response to the initial perturbation to coordinate multicellular transcriptomic responses. Additionally, ReCoN predicted distinct fibroblast states emerging in different microenvironments reconstructed from spatial data. ReCoN provides an interpretable modeling framework for multicellular systems that allows for the simulation of perturbations, including the assessment of the cellular selectivity of these treatments *in vivo*. Ultimately, it can help design patient-specific molecular therapies.

**Graphical abstract:** 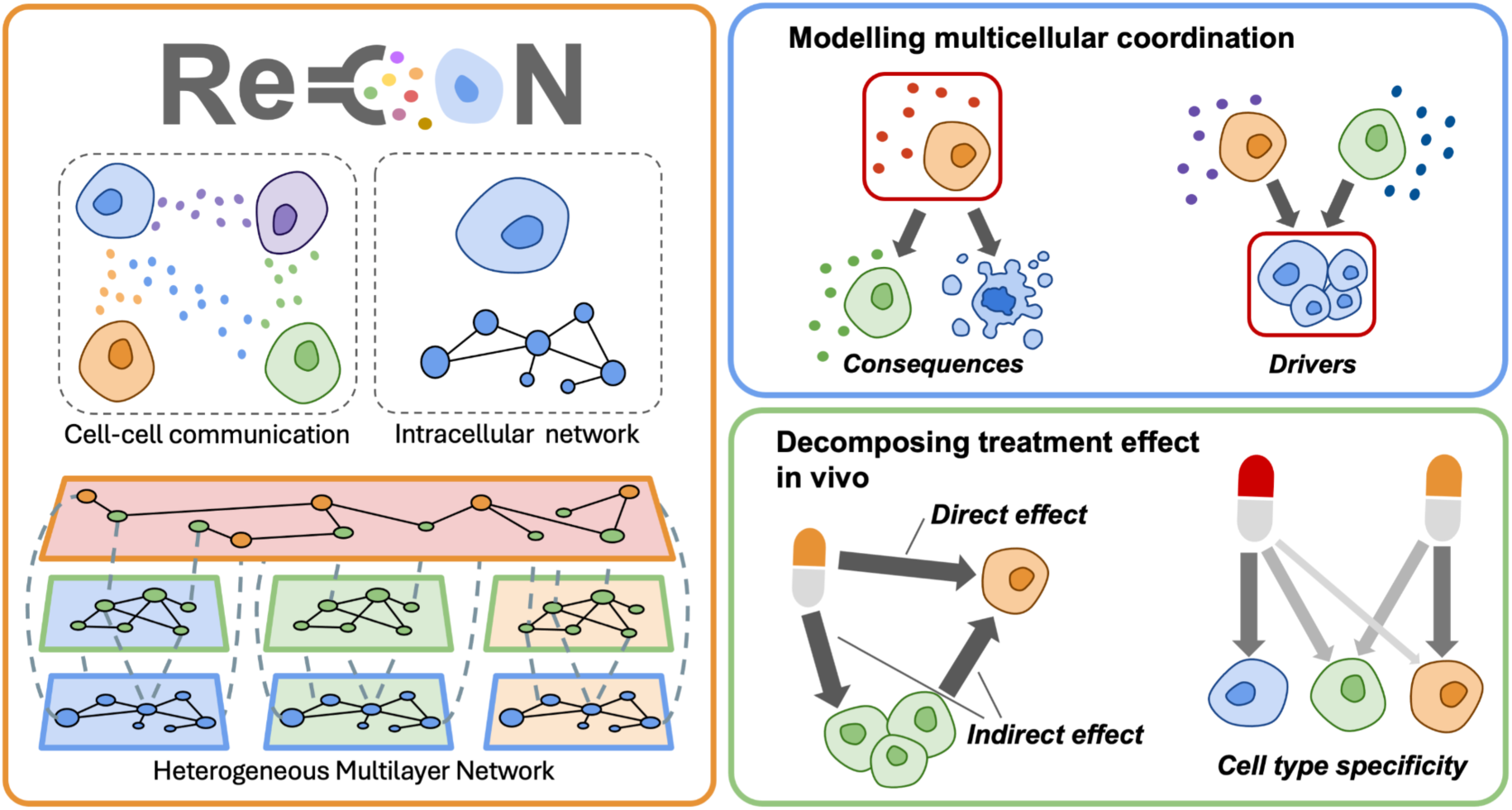

## 1. Introduction and background

A cell’s identity depends on internal molecular mechanisms and its environment^1–3^. In multicellular organisms, the environment includes surrounding cells that exchange molecular signals, modulating how cells respond to perturbations. The complex coordination that emerges, coupled with intrinsic cell diversity, shapes tissue functions via multicellular programs. A molecular cue influences cells both directly through receptor-mediated pathways and indirectly, via effects on other cells that in turn send secondary signals^4–6^. These indirect routes can significantly shape the final transcriptional response, and capturing both effects is essential for accurate modeling of biological responses.

Since the emergence of single-cell technologies, cell coordination has largely been studied through the inference of cell-cell communication via ligand–receptor expression analysis^7^. These approaches have been effective in identifying signals conveying information between cells. Some methods further extend this framework by predicting downstream intracellular targets of receptor activation^8–11^, partially bridging the gap between extracellular signaling and gene regulation. However, cell–cell communication analyses typically remain limited to pairwise interactions and do not model the complete multicellular coordination occurring in complex tissues.

More recently, tissue-centric approaches have been developed to characterize coordinated gene expression across multiple cell populations^12,13^. In particular, multicellular program analyses^14–17^ decompose multi-sample single-cell or spatial transcriptomic data into latent factors representing coordinated gene expression patterns across distinct cell types. While these approaches effectively describe *what* tissue-wide programs are coordinated, they generally offer limited insight into the molecular mechanisms that establish or regulate these programs.

Despite their ability to identify coordinated tissue-wide programs, multicellular program analyses typically offer limited insight into the underlying molecular mechanisms that orchestrate these programs. On the other hand, cell–cell communication inference tools can identify molecular signals between pairs of cells, but do not model how these signals dynamically propagate to organise complete multicellular programs. These limitations highlight the need for methodological frameworks that combine the strengths of both approaches to better understand how multicellular behaviours emerge in complex tissues.

We here present ReCoN (REconstruction of multicellular COordination Networks from single-cell data), a computational framework designed to model multicellular coordination and responses to perturbation by integrating intracellular regulation and cell-cell communication. ReCoN combines tissue- or cell type-specific intracellular regulatory networks with inferred ligand–receptor communication graphs in a multilayer network. This explicit modelling allows tracing how molecular signals spread through interacting cell types, based on random walk with restart explorations. ReCoN also formulates the direct and indirect effects across tissues separately, allowing for weighing their individual contributions.

We demonstrate that ReCoN outperforms existing baselines at predicting the effect of different ligands in both observational and interventional contexts, leveraging the indirect effects described above. We first predicted the cell type responses to different cytokines from the Immune Dictionary, an in vivo perturbation dataset, evaluating the contribution of each ReCoN’s component. We also applied ReCoN to a more complex context, retrieving genes relevant for heart failure from a combination of ligands involved in their coordination across cardiac cell types. ReCoN was then applied to reconstruct three post-cardiac infarction microenvironments, identified from spatial transcriptomic data. Distinct and coherent fibroblast phenotypes were predicted from each microenvironment model. We finally explored the regulatory mechanisms behind cardiac fibrosis and heart failure. We identified both molecules and larger biological programs across cardiac cell types interacting with fibroblasts, before and after fibrosis activation.

## 2. Results

### 2.1. ReCoN: a new method to extract molecular mechanisms in multicellular systems

Understanding how cells coordinate distinct molecular programs requires bridging intracellular regulation with intercellular signaling. We present here ReCoN (Regulatory mechanism and Cell Communication Network), a network-based approach integrating both scales and modelling multicellular molecular coordination.

#### 2.1.1. ReCoN: a heterogeneous multilayer network framework

We developed ReCoN, a new method for modeling multicellular systems and exploring the consequences of multicellular coordination on molecular perturbations, leveraging single-cell multi-omics data and prior knowledge. Through its mechanistic approach, ReCoN identifies the direct effect of signal transduction and the indirect effects of the environment through cell-cell communication.

To model multicellular coordination and its consequences, ReCoN starts by creating a network for each cell type of interest. Called cell type multilayers, they are based on a heterogeneous multilayer structure (HMLN), which allows the integration of multiple types of biological networks^18–22^ (Figure 1a). The cell type multilayers include two layers connected by the receptor-gene bipartite network: a gene regulatory layer and a receptor layer. The gene regulatory layer contains transcription factors and their target genes, inferred from scRNA-seq or single-cell multi-omics data^23^. This layer can be inferred with ReCoN directly or another GRN method. The receptor layer represents membrane proteins, and can optionally include receptor–receptor interactions, such as dimerization. In order to propagate signals in-between them, these two layers need to be connected. To do so, we inferred the transcriptional impact of cellular receptors, and we provide receptor-gene links for both mouse and human that can be used as a bipartite network linking these molecular levels. These links have been obtained by decomposing the ligand-gene links from Nichenet^24^ into ligand-receptor and receptor-gene links (see Methods). Each cell type multilayer is independent, which allows for the representation of cell–type–specific regulatory mechanisms (Supplementary Notes 1).

**Figure 1.**
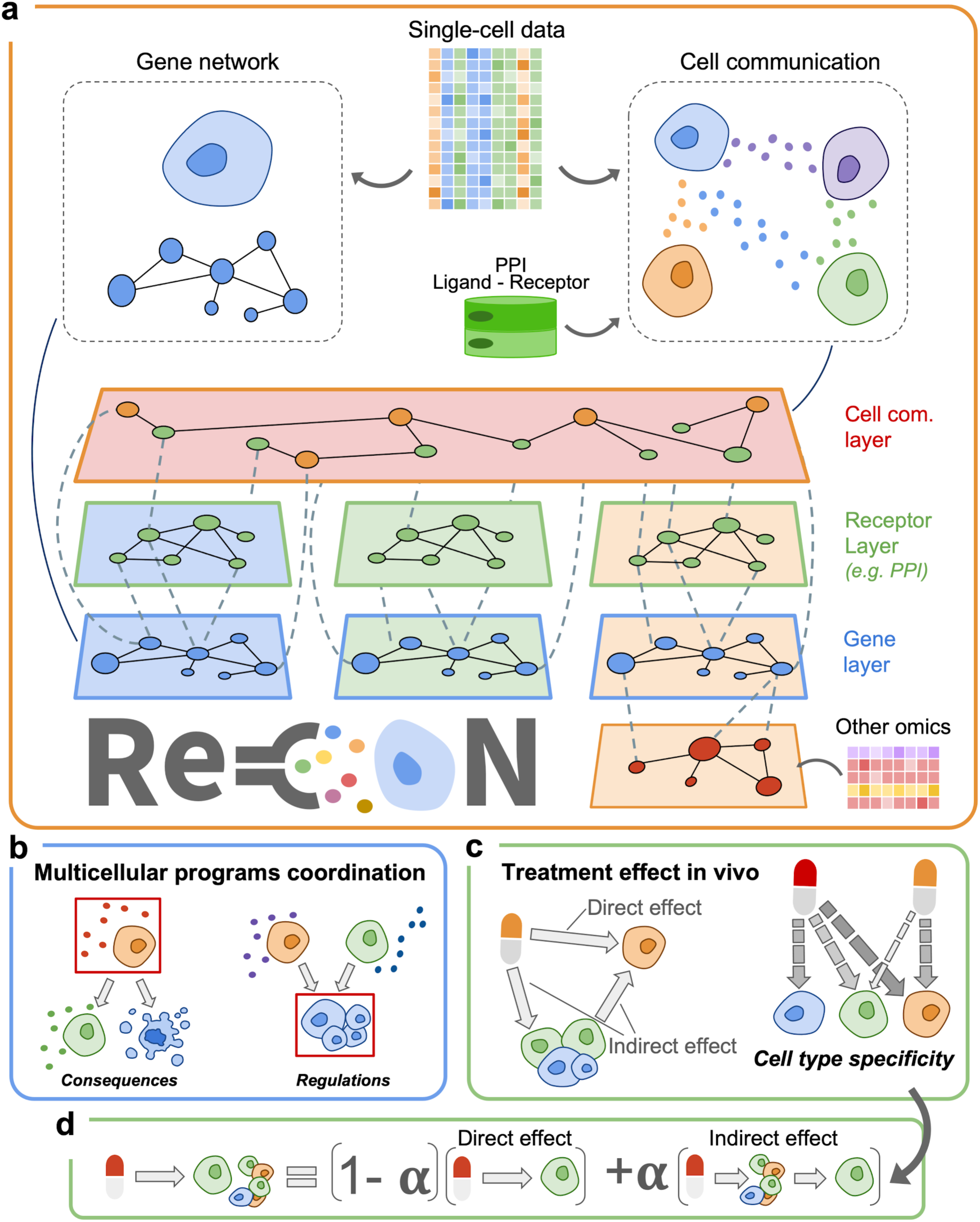
An overview of the framework and applications of ReCoN. **a)** General HMLN structure used in ReCoN to connect intracellular network layers and a cell-cell communication layer. Each cell can contain a different number of layers, illustrated by the asymmetric structure presented. **b)** ReCoN can be used to infer both the upstream driver and downstream target cross-cellular signals of a profile or pathway of interest (grey frame). **c)** ReCoN decomposes the effect of a molecule into a direct and an indirect effect. The direct effect on a cell corresponds to the initial signal transduction, while the indirect one occurs in response to the surrounding cells’ signals. **d)** The molecular changes in a cell type after a treatment in vivo result from both the direct and indirect effects. Their contributions are weighted using *α*, and the impact of each cell type *i* is also modulated by *B_i_*_→*j*_coefficients.

The cell type multilayers are then connected into a large multicellular network by an external cell-cell communication layer, inferred from scRNA-seq data and a prior database of ligand-receptor bindings. The nodes of this network are ligand/receptor-cell type pairs, indicating from/to which cell type multilayers they are connected. To allow transitions between intercellular and intracellular interactions, the cell-cell communication layer is then connected to the cell type multilayers through two bipartite networks (see Methods). Trivially, the ligand-cell type nodes are linked to their corresponding genes in the gene regulatory layers, and the receptor-cell type nodes are connected to the matching receptors in the receptor layers.

Formally, this network *N* = (*V*, *V_m_*, *E_m_*, *L*), *m* = 1, …, *M*, is composed of *M* layers containing different *V* nodes and *V_m_* their multiple representations across the layers, or node-layer tuples. The edges between node-layer tuples are encoded in *E_m_* ⊂ *V_m_✕ V_m_*^25,26^. The properties of the layers are specified in *L*. Since each graph can have its own properties, it offers a great general framework to connect different types of molecular interactions and different compartments (intracellular and extracellular spaces). By default, each layer is undirected and weighted. However, the multilayer framework of ReCoN allows for the integration of networks with different properties, as long as all edge weights remain positive.

Once the multicellular network is created, ReCoN aims to find the regulators and downstream targets of a molecular signal (Figure 1b). ReCoN measures signal propagation through random walk with restart (RWR)^18^ (see Methods). This algorithm begins with one molecule of interest, such as a gene/receptor, or several molecules, such as a pathway, and computes a probability distribution over the entire network’s nodes. This distribution reflects the strength of each node’s connection to the input. It can be interpreted as the multicellular effect of a perturbation (or multicellular regulators, depending on the direction of exploration). The random walk balances steps within and between layers using a transition matrix, exploring the complete multicellular network. General transition matrices define the types of exploration for discovering regulators or downstream targets (see Methods).

In the case of extracellular perturbation, ReCoN can reconstruct the secondary (later on called indirect) effect of perturbations. In this case, a first random walk is then computed solely allowing receptor-binding and intracellular transitions. Secondly, from the ligands reached in each target cell type, another random walk, allowing intercellular transitions, is ran to identify its secondary signals.

ReCoN and its dependencies are working directly on AnnData objects, facilitating their integration downstream of other scVerse packages. To infer the gene regulatory network, ReCoN embeds a Python version of HuMMuS^21^, which considers interactions between DNA regions by also creating an HMLN. The code of ReCoN is accessible at https://github.com/cantinilab/ReCoN, and its documentation can be found at https://recon.readthedocs.io.

#### 2.1.2. ReCoN defines the direct and indirect effects of a perturbation

ReCoN distinguishes between two types of regulatory effects: direct and indirect (see Figure 1c). The direct one captures the influence of the initial signal on each cell type via their respective intracellular signal transductions. The indirect effect corresponds to the impact of intercellular interactions. Surrounding cells are also stimulated by any molecule (e.g., cytokine) and, in turn, release ligands or signals. Such indirect signaling can then affect the target cell through cell–cell communication (CCC).

This distinction is formalized through two types of random walk with restart (RWR) on the heterogeneous multilayer network (see Figure 1d). The key difference between them lies in the ability to transition from the intracellular layers to the cell-cell communication layer. A parameter *γ* controls this transition probability, effectively tuning the influence a cell type has on its surrounding environment.

Formally, the direct effect (*M*_*D*_) is computed by running an RWR on the full network *G*, starting from the input molecular profile of interest *M_0_*. This input is a vector encoding the importance of each input node, as the probability of restarting the RWR from them. In this RWR, transitions from intracellular to intercellular layers are blocked ( *γ* = 0), to capture only the intracellular effects of the input, as shown in (*1*):

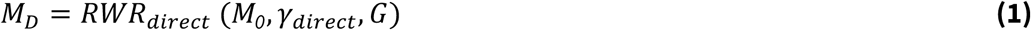

The result of this step is a vector *M_D_*, representing the distribution of the direct effect across the network. This vector also serves as the input for computing the indirect effects, which model the downstream signaling between cells.

Indirect effects are calculated using additional RWRs that allow transitions between intracellular and communication layers. By default, transitions between these layers and staying within a layer are equiprobable (*γ_indirect_* = *0*.*5*). For each cell type *i*, ReCoN identifies the genes coding for ligands that act through direct effect, *mask_ligands i_* (*M_D_*). A RWR is then run from these ligands, capturing the impact of that cell type over the others. These indirect effects are weighted by cell-type-specific influence vectors *B_i_*, and the results are aggregated across all emitter cell types. *B_i_* typically corresponds to the cell-type proportion, but can be adapted to encode the cell-type proximities or specific cooperation scores.

Finally, the overall system response *M_T_* is obtained by a convex combination of the direct effect and the indirect effects, as shown in (*2*):

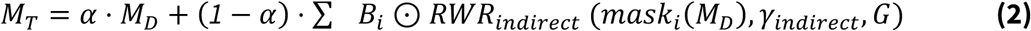

In conclusion, ReCoN aims to bridge intracellular regulation and cell-cell communication to predict the effect of molecular perturbations. By separating the direct from the indirect effects, ReCoN allows weighting their respective contribution to multicellular system perturbations. ReCoN’s HMLN structure leverages both prior knowledge and contextualised networks from single-cell multi-omics data to find new and specific interactions. Through its modular aspect, cell types and omics can be easily added to match the hypothesis and environment of interest. It also allowed us to measure the contribution of the different layers of the ReCoN network. ReCoN exploration can take several molecules as input, modelling their joint effect.

### 2.2. ReCoN predicts multicellular responses to cytokine treatments in murine lymph nodes

In vivo tissue studies provide a biologically relevant setting to evaluate ReCoN, as they reflect the coordinated interactions among diverse cell types. We therefore measured ReCoN performance and the individual contribution of each of its components on the Immune Dictionary dataset, which profiles murine lymph node responses to a panel of cytokine treatments administered in vivo^27^. Transcriptomes from 17 immune cell types were sequenced, offering a rich opportunity to study cell type responses coordination within a complex tissue (see Figure 2a).

**Figure 2.**
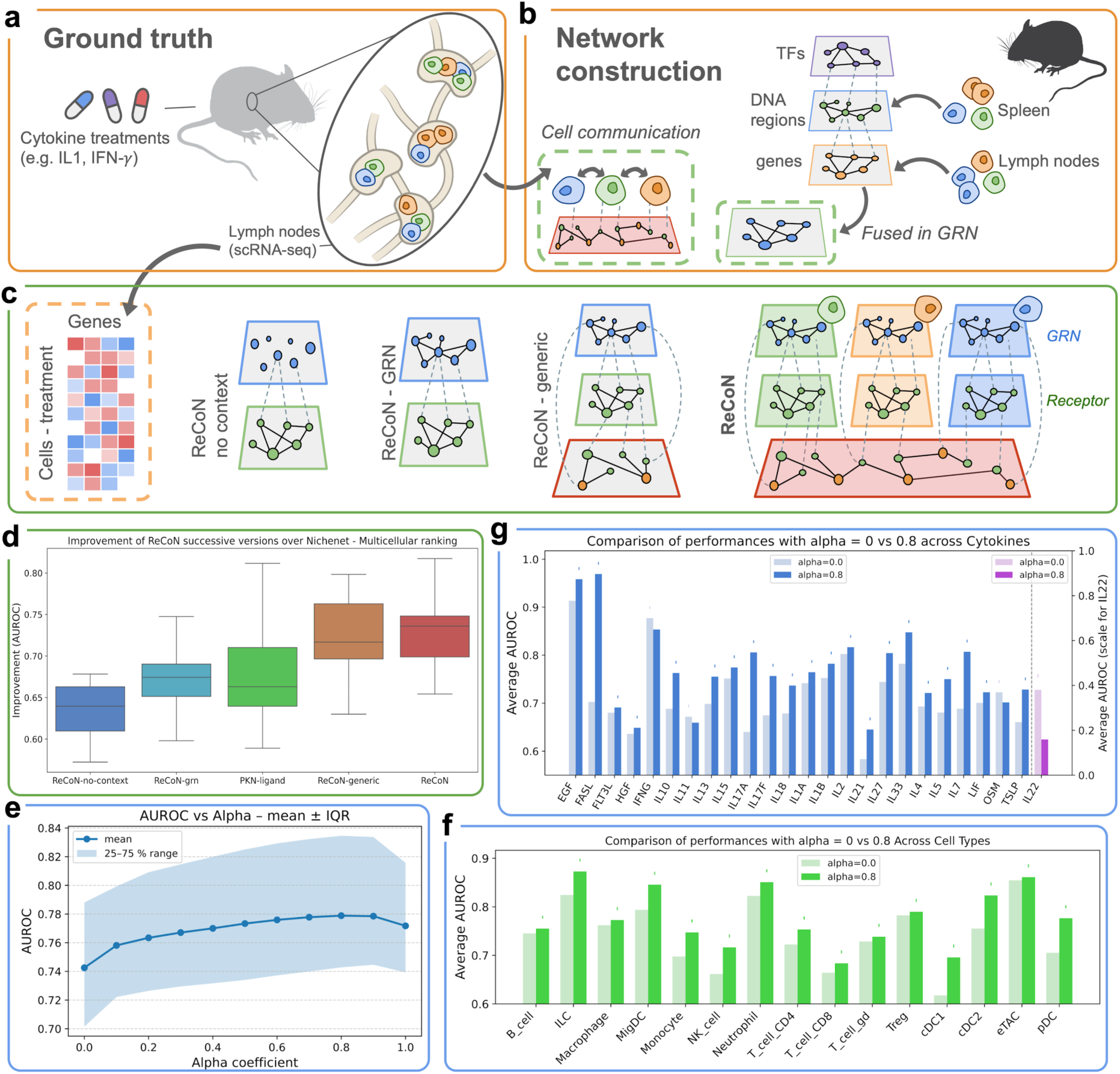
ReCoN recovers cell type responses from different cytokine treatments, leveraging cross-cellular effects. **a)** Schematic representation of the Immune Dictionary dataset used in performance evaluation. **b)** Illustration of the data used to build ReCoN’s networks. **c)** Schematic view of the successively enriched networks used in d. **d)** Cytokines AUROC distributions for the four successively enriched versions of ReCoN (dark blue, light blue, orange, and red) and the ligand-PKN model (green). **e)** Cytokine-cell type pairs AUROCs distributions for different α (indicating the contribution of the indirect effect) in ReCoN. **f)** Average AUROCs for each perturbation with α = 0.0 (in light green, only direct effect) and α = 0.8 (blue, strong indirect effect); best performance is indicated by a dot. **g)** Average AUROCs for each cell type with α = 0.0 (in light green, only direct effect) and α = 0.8 (blue, strong indirect effect); best performance is indicated by a dot. *The mice and lymph node illustrations in b come from the NIH BioArt Source library – bioart.niaid.nih.gov/bioart (20, 589)*.

Since this dataset only contains scRNA-seq data and very few unperturbed cells, we used external data to build the cell type multilayers (see Figure 2b). The gene regulatory network was inferred from both a lymph node scRNA-seq dataset and a splenic scATAC-seq dataset from the same mouse inbred strain. The cell-cell communication network was inferred from the Immune Dictionary cells, treated with phosphate-buffered saline (PBS), used as a negative control.

#### 2.2.1. Gene regulation and cell-cell communication networks are both informative

To assess the contribution of each context-specific information in ReCoN, we benchmarked four variations of the ReCoN methodology (see Figure 2c). They correspond to two unicellular versions, ReCoN-no-context and ReCoN-grn, and two multicellular ones, ReCoN-generic and ReCoN. Altogether, these models illustrate the progressive integration of regulatory and communication layers in ReCoN. We also compared ReCoN to the ligand–gene prior knowledge network (ligand-PKN) from NicheNet^24,28^, which has been used to derive the receptor-gene bipartite for ReCoN’s models (see Methods).

The first ReCoN variation tested, ReCoN-no-context, includes only receptor–gene links derived from the ligand-PKN. It thus ignores GRNs and CCC to isolate the contribution of the context-independent receptor-gene prior knowledge network (PKN). This minimal configuration also serves to assess how much predictive signal was lost from the ligand-PKN. The second variation, ReCoN-grn, includes a context-specific GRN built from single-cell RNA-seq data from lymph nodes and ATAC-seq data from the spleen. The GRN is shared across all cell types, and no CCC network is included. This model represents the direct effects of cytokine perturbations while adding tissue-specific regulatory context. ReCoN-generic adds a generic CCC network on top of the GRN, connecting all possible ligand–receptor pairs irrespective of cell type expression. It highlights the contribution of indirect effects arising from cell-cell communication without personalized interaction specificity. Finally, the full ReCoN model incorporates both context-specific GRNs and a personalized CCC network based on cell-type-specific ligand and receptor expression. This model captures how different cell types contribute to each other’s regulation through the expression of ligands and receptors, reflecting multicellular tissue organization. In both ReCoN-genetic and ReCoN, we can modulate the importance of cell-cell communication and its indirect effects through the coefficient α (see 2.1). We show here the results with α = 0.8, presenting the best performance, before exploring the impact of this parameter in the next session.

We first evaluate the models’ ability to predict the transcriptomic changes of each cell type separately, through the individual cell type gene rankings (see Supplementary Figure 1). We then verify the models’ ability to combine cell type scores across the tissue through a multicellular ranking (see Figure 2d). This multicellular evaluation tests whether a model can prioritize strongly perturbed genes from different cell types, without bias toward one of them. It thus considers its capacity to combine their different score distributions, even if several cell types present very different response profiles and intensities.

The performances of the models are reported as area under the receiver operating characteristic curve (AUROC) values. As expected, ReCoN-no-context presented the worst performances in both the individual cell type and multicellular predictions. It underperformed compared to the ligand-PKN model, with associated Mann-Whitney U tests P-values of 3.32e-2 and 6.53e-2, respectively. This illustrates the initial loss of information when inferring the receptor-gene links from the ligand-PKN. The ReCoN-grn outperformed ReCoN-no-context on both metrics (Mann-Whitney U test P-values of 8.13e-4 and 9.32e-3). This significant improvement demonstrates the contribution of the personalized GRN in predicting in-tissue cell-type perturbations. It additionally performed slightly better than the ligand-PKN model, thus compensating for the previous information loss. Finally, the ReCoN-generic and ReCoN models demonstrated the best performances, significantly outperforming the ReCoN-grn model on both metrics (ReCoN-generic P-values of 3.88e-6 and 4.61e-3, and ReCoN P-values of 5.02e-7 and 3.83e-3). This highlights the necessity of considering the indirect effect through cell-cell communication to predict cell perturbation responses accurately. The full ReCoN, which additionally considers the cell type specificities, showed a modest performance increase on the individual cell type evaluation, and outperformed ReCoN-generic more clearly on the multicellular scores. Although the average AUROC improved from 0.71 to 0.73 with the personalized ReCoN model, the associated P-value of 7.86e-1 indicates that this difference is not statistically significant. Despite a modest statistical improvement, the complete model of ReCoN is particularly useful in the context of multicellular cooperation since it allows to explore cell-type-specific responses.

Overall, integrating context-specific GRNs and CCC networks enhances the prediction of cytokine-induced gene expression changes. The inclusion of CCC information, in particular, underscores the significance of indirect effects mediated through intercellular signaling. These findings highlight the importance of considering both intrinsic regulatory mechanisms and extrinsic communication cues for accurately modeling multicellular responses. It is expected to observe common behaviors in-between cell-type, that the GRN and the generic CCC network already contribute captures. However, being able to predict cell-type specificities while conserving the same performances is here a great benefit of the complete model. Importantly, the benefit of personalizing CCC is likely underestimated here, as its inference from spatially agnostic single-cell transcriptomics-based methods typically lead to many false positives^8^.

#### 2.2.2. Indirect effects and cell-cell communication explain most of the cytokine treatment responses

We additionally assessed the relative influence of the cellular environment versus the direct signal transduction. We evaluated ReCoN’s performance across a range of alpha (α) values, which balance the direct and indirect effects contributions. Overall, the best response predictions averaged per cell type were achieved at α = 0.8, indicating a dominant contribution of indirect signaling via cell–cell communication (see Figure 2e). Additionally, the mean AUROC across all cytokine–cell type pairs was 0.76 at α = 0.8, compared to 0.72 at α = 0 (direct effect only), with a Mann–Whitney U test p-value of 1.07 × 10⁻⁵. Consequently, α = 0.8 was selected as the default parameter in subsequent analyses of ReCoN-generic and personalized ReCoN models.

We further investigated the relative impact of indirect effects per cell type. We first measured the optimal α for each cell type. For all, the best performances were associated with an α value superior to 0.5 (see Supplementary Table 1). We furthermore quantified the predictive power gained by considering cellular coordination (α = 0.8) over the direct effect alone (α = 0) (see Figure 2f). Dendritic cells showed the largest gains at α = 0.8, with relative improvements of 12.6%, 10.1%, and 9.0% for pDC, cDC1, and cDC2, respectively. This is consistent with their known roles in orchestrating immune responses and recruiting other cell types^29,30^. Dendritic cells also exhibit high functional heterogeneity and adaptation capacity, notably in response to tumor-specific environments^31^. In contrast, eTACs and Tregs showed the smallest improvements, consistent with their high degree of functional specialization. Both are known for their role in immunosuppression and the inhibition of T cell activation^32,33^. Notably, eTAC-mediated CD4⁺ T cell inactivation has been shown to occur independently of both Tregs and innate inflammatory stimuli^32^.

This illustrates the broad and systematic contribution of indirect effects across the studied cell types. The amplitude of this contribution varied between cell types, reflecting differences in their sensitivity to the environment. These variations were consistent with known functional roles, with some cell types showing greater capacity to adapt their responses to contextual signals.

We next examined whether the balance between direct and indirect effects varied between cytokines. Some molecules may exert stronger cell-intrinsic effects, particularly those that drive decisive processes such as apoptosis or proliferation. Among the 22 cytokines tested, 17 (77%) achieved optimal performance with α > 0.6 (see Supplementary Table 2), indicating that indirect effects were generally dominant. IFN-γ and FLT3L showed better predictions at α = 0.2, IL11 at α = 0.2, and OSM and IL22 at α = 0, suggestive of a stronger direct component. IFN-γ performance was an exception, declining markedly at high α values, which is consistent with its well-documented cell-intrinsic pro-apoptotic activity^34^. FLT3L showed only modest variation, and the indirect effect alone still outperformed the direct one. FLT3L is primarily involved in immune cell proliferation and migration, amplifying and modulating context-dependent signaling^35^. IL11 and OSM also exhibited minimal change across α values, while IL22 remained poorly predicted regardless of α. Due to very few genes being differentially expressed, only cDC1 was conserved and evaluated for IL22, which could also explain the poor performance (see Supplementary Figure 2). When excluding IL22, all cytokines had a relative improvement between -2% and +38%, with an average of 8% using α = 0.8 (see Figure 2g).

Overall, cytokine responses reflect a balance between direct and indirect regulation. While indirect signaling predominates across most treatments, the relative contribution of direct effects varies among cytokines. Further investigation of these variations across a broader range of cytokines could enable more precise modeling of the direct–indirect effect balance tailored to each molecule.

### 2.3. ReCoN predicts heart failure transcriptomic changes from potential key regulators

ReCoN can also model the impact of more complex molecular perturbations. We applied and evaluated ReCoN for retrieving the coordinated multicellular response occurring in heart failure (HF). In ReHeat2, a meta-analysis of human HF transcriptomic data^36^, the authors identified widespread shifts in cell type composition and gene expression, including a multicellular program separating HF from non-failing hearts (NFH) (see Figure 3a). This program was associated with ligand–cell type pairs potentially driving disease progression. We focused on these ligand scores to test whether ReCoN could recover their joint regulatory impact. Therefore, we used these ligands as inputs, weighting their influence based on their scores, to model their combined regulatory impact across cell types.

**Figure 3.**
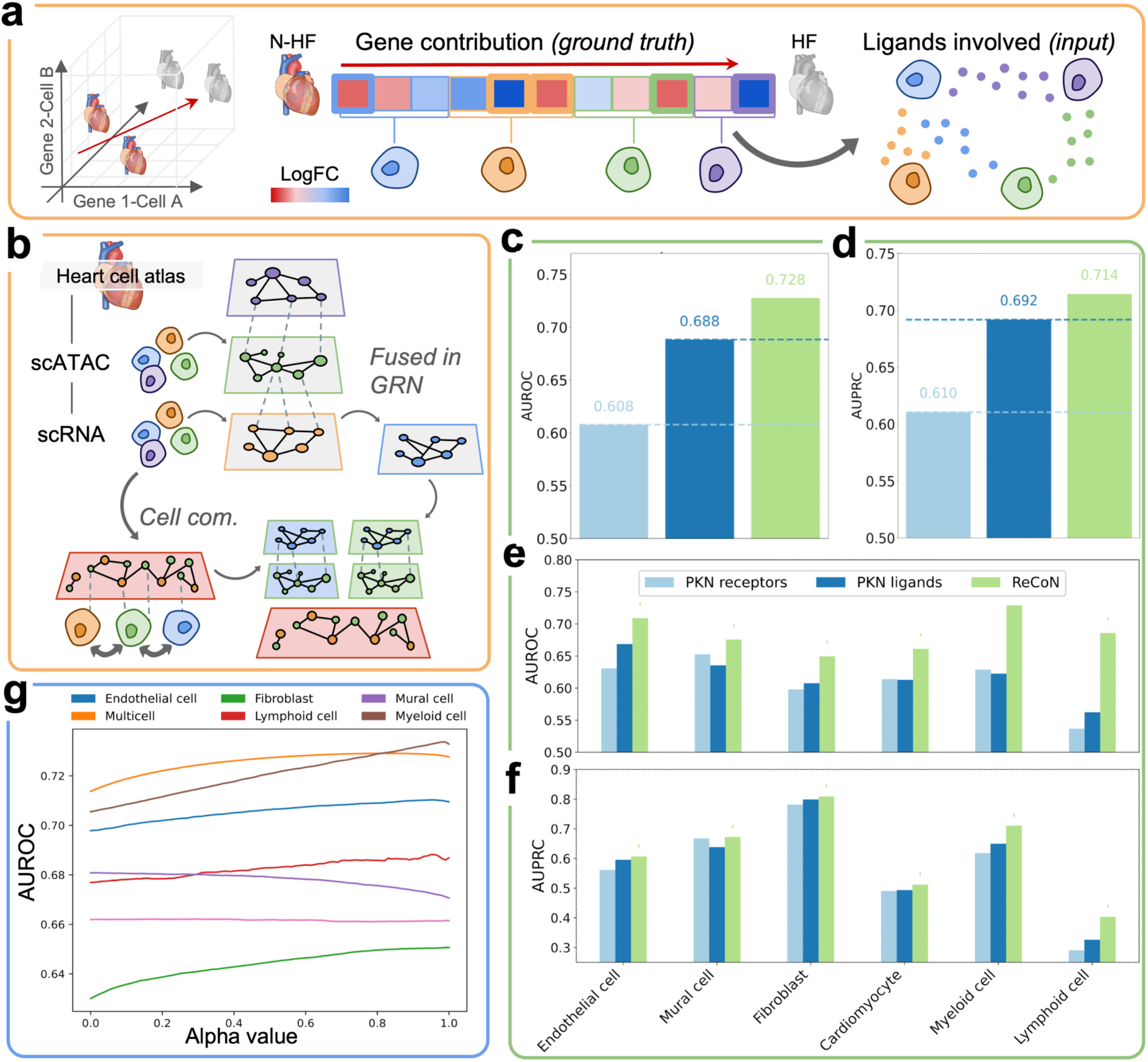
ReCoN retrieves the multicellular transcriptomic response to ligands involved in Heart Failure. **a)** Schematic overview of the heart failure data used as ground truth for model evaluation. **b)** Schematic representation of the ventricular heart model reconstructed with ReCoN. **c)** AUROC for predicting all differentially expressed genes in the multicellular heart model (i.e., global ranking across all cell types), comparing ReCoN (green) to two prior knowledge–based models: PKN-ligand (light blue) and PKN-receptor (dark blue). **d)** AUPRC for the same multicellular prediction task as in panel c. **e)** AUROC for predicting differentially expressed genes in individual cell types, comparing ReCoN to PKN-based baselines. **f)** AUPRC for the same gene prediction task in individual cell types and the same models as in panel c. **g)** AUROC as a function of α (the weight of indirect effects) for individual cell types and global multicellular rankings. *The heart illustrations in a and b come from the NIH BioArt Source library – bioart.niaid.nih.gov/bioart (228)*.

The underlying network was built from a multi-omics dataset of human left ventricle samples^37^, including six cell types: cardiomyocytes, fibroblasts, myeloid cells, lymphoid cells, mural cells, and endothelial cells (see Figure 3b, Methods). We compared ReCoN’s performance to two reference models derived from NicheNet: the original ligand–target gene network (ligand-PKN) and a receptor-centric version (receptor-PKN), which links receptors to target genes via inferred ligand–receptor interactions. AUROC and AUPR were used to assess model performance in both multicellular and per–cell-type gene rankings.

In the multicellular ranking, which assesses how well models capture effects across multiple cell types in a comparable manner, ReCoN outperformed both PKN-based approaches in AUROC (see Figure 3c) and AUPR (see Figure 3d). This ability to jointly evaluate regulatory influence across cell types is essential for identifying selective interventions that affect target populations while sparing others. ReCoN also outperformed the baseline models in the per–cell-type rankings, achieving higher AUROCs (see Figure 3e) and AUPRCs (see Figure 3f) in all six cell types. The AUROC improvement over ligand-PKN ranged from 22% in lymphoid cells to 6.0% in endothelial cells, while the AUPRC improvement ranged from 24% in lymphoid cells again to 1.3% in fibroblasts. Lymphoid cells display especially distinct receptor expression profiles, making them benefit particularly from ReCoN’s incorporation of cell-specific signaling context.

We next assessed how ReCoN’s performance varied with the relative contribution of the direct and indirect effects. In this setting, the best multicellular and per–cell-type rankings were achieved with α values above 0.5 for all cell types, except mural cells (see Figure 3g). The alpha associated with the best performance on the multicellular ranking was 0.82, close to the 0.8 identified from the Immune Dictionary dataset. It again emphasizes the contribution of indirect signaling and validates this default value by two different systems in mouse and human.

ReCoN consistently outperformed the PKN-based models across both multicellular and per–cell-type evaluations. Unlike the Immune Dictionary, which captures interventional dynamics with clear temporal resolution, the heart failure analysis is observational and lacks information on signaling chronology. As a result, distinguishing direct from indirect effects is less straightforward: disease-associated ligands may actually initiate cell-intrinsic responses or act downstream to modulate multicellular coordination. Nevertheless, the contribution of indirect effects underscores again the importance of considering cell type interactions to understand their responses even in complex, non-interventional contexts.

### 2.4. Modeling the microenvironment impact on fibroblast phenotype in heart infarction recovery

ReCoN can also be applied to model and contrast different microenvironment patterns of the same organ. Spatial transcriptomic methods offer the opportunity to identify local cell type neighborhoods that can all react differently and express different signals.

In heart infarction, notably, since the lesion can be highly localised, the cardiac tissue recovers very heterogeneously. Some regions stay highly functional, keeping a high proportion of myocardiocytes. In contrast, others have an inflammatory profile and an increased number of immune cells, or a fibrotic signature and differentiation of fibroblasts into myofibroblasts. These regions will recover from tissue damage differently, each involving a particular interplay of the cardiac cell types. Notably, fibroblasts have a major role across the three contexts^38,39^, identifiable by distinct cell states.

In this showcase, we predict fibroblast subtypes profiles in three spatial contexts – inflammatory, fibrotic, and functional – using only information from the other cardiac cell types. We aim to verify that the surrounding cardiac cell types can explain the distinct fibroblast profiles found in these spatial contexts.

We used Visium 10X spatially resolved transcriptomic data from a large heart infarction study^39^ to identify the three spatial contexts and build the corresponding models. We selected a slide containing all three spatial contexts of interest, including the ischaemic zone located near the lesion (see Figure 4a). For each spatial spot, we inferred both the cellular composition and the gene expression attributed to each cell type with cell2location. The gene expression attributed to each cell type can be compared to spot-specific pseudo-cells and used here in the same way as single-cell profiles (see Figure 4b, Methods). It was used both to infer the cell network and the average profile of each cell type, interpreted as the environment input signal for the fibroblast profile predictions. The GRN representing intracellular mechanisms was the one computed in the Heart Failure showcase (see Methods).

**Figure 4.**
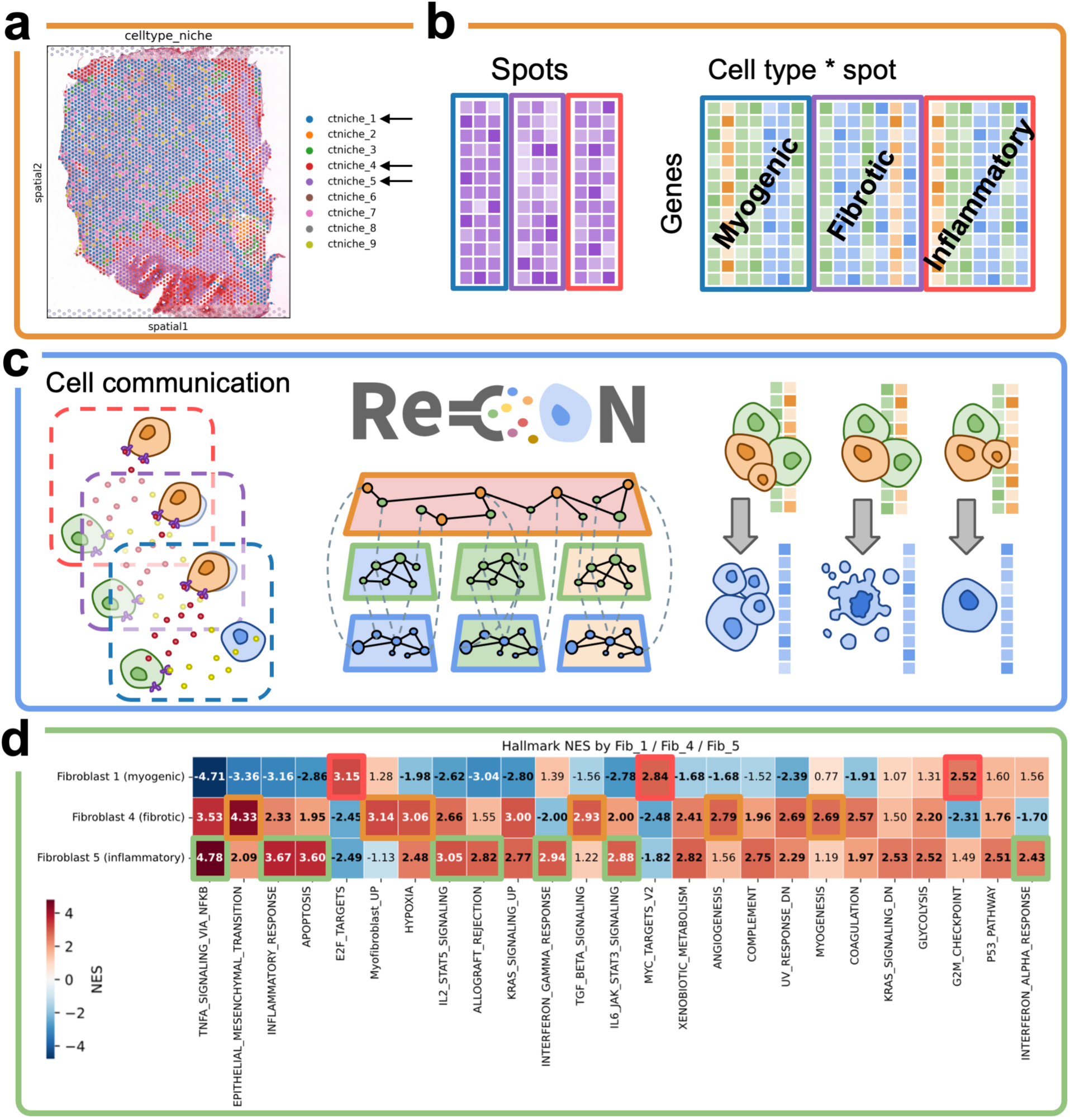
ReCoN predicts different fibroblast specializations depending on the cardiac microenvironment. **a)** Spatial mapping of compositional niches on an ischaemic zone sample post myocardial infarction. **b)** Schematic view of the deconvolution process and cell type-specific count inference from the spatial niches. **c)** Workflow used to infer the different fibroblast states from the environment only. We first use the predicted spot*cell type count of each type of niche to infer cell-cell communication networks. A ReCoN network is built around each of them. Then, using the average gene expression for each cardiac cell type but the fibroblast of each spatial context, we explore our ReCoN model and obtain the predicted profile of the fibroblast. **d)** Gene sets enrichments of general Hallmarks and myofibroblast markers by the predicted fibroblast profiles in the three studied spatial contexts: myogenic, fibrotic, and inflammatory. Pathways of interest for each context are surrounded.

The average profile of the other cardiac cell types constitutes a large multicellular vector that was used as a seed for ReCoN’s RWR exploration. We predicted the downstream profiles of fibroblasts from these three multicellular environments and compared them using the log-ratio of these predictions against the mean of the other context groups (see Figure 4c). We then enriched general molecular hallmarks and markers of characteristic fibroblast states, such as myofibroblasts, to identify transcriptomic differences.

ReCoN was able to recover different pathways characteristic of each region (see Figure 4d). In the fibrotic region only, the fibroblasts significantly enriched (p-value < 0.05) several gene sets highly related to fibrosis, such as TGF beta signaling, the myofibroblast marker set, and the myogenesis hallmark. Myofibroblast is indeed a cell state highly associated with wound healing and fibrotic tissues^40^. In addition, the epithelial-mesenchymal transition (EMT), angiogenesis, and hypoxia hallmarks were significantly enriched in both the fibrotic region and, at a lower level, in the inflammatory region, highlighting common adaptation of fibroblasts to damaged tissue microenvironments. In contrast, the inflammatory region fibroblasts were the only ones that enriched positively and significantly the IFN-*γ* and IFN-*α* signaling pathways, which were negatively enriched in the fibrotic region. Other inflammation-related gene sets, such as inflammatory response hallmark, TNF-*α* signaling, IL2 and IL6 signaling, and apoptosis, were positively enriched in the inflammatory region and negatively enriched in the fibroblast of the myogenic region. Here again, the fibrotic context also enriched these pathways at a lower level, illustrating closer proximity between both the inflammatory and the fibrotic contexts than with the myogenic one. Fibroblasts from the myogenic region positively enriched fewer gene sets significantly, which included notably the target gene sets of very generic TFs, E2F, and MYC.

Using only their respective microenvironment, ReCoN was able to predict the difference between several fibroblast profiles. We evaluated our predictions by enriching gene sets coherent with the expected transcriptome of the fibroblasts in inflammatory, fibrotic, and myogenic regions. It demonstrates the role of neighboring cells in determining a cell’s fate. Altogether, these results suggest that ReCoN can be used to reconstruct the effect of different cellular environments, notably to explain the distinct cell states observed for a cell type of interest. More generally, we aimed in this showcase to demonstrate ReCoN’s potential in reconstructing different patients, tissues, and microenvironments to predict context-specific molecular treatments.

### 2.5. Retrieving cell type-specific drivers of cardiac fibrosis in different molecular levels

Heart failure arises from the co-occurrence of multiple physiological and cellular modifications that disrupt cardiac structure and function. Cardiac fibrosis, a hallmark of this pathological remodeling, is driven primarily by the activation of cardiac fibroblasts and their transition to myofibroblasts, which secrete excessive amounts of extracellular matrix (ECM) proteins. This abnormal ECM accumulation stiffens the myocardium, impairs contractility, and promotes arrhythmias. We investigated here the specific coordination of major cardiac cell types with cardiac fibroblast and ECM production, in order to identify potential ligands and intracellular regulators of fibrosis.

In contrast with the previous showcases, we focused here on predicting the regulators of molecular perturbations. We used ReCoN to identify the molecules regulating the production of ECM proteins in the HF model (see Figure 5a). First, we combined 10 gene sets from mSigDB related to ECM production^41^, excluding the ones related to cancer propagation, to form a set of genes related to fibrosis. Starting from this gene set in the fibroblast network, we used ReCoN to predict potential upstream TF, receptors, and ligands regulating fibrosis activation. To do so, the RWR exploration is set up to only explore upstream interactions in-between molecular layers (see Methods).

**Figure 5.**
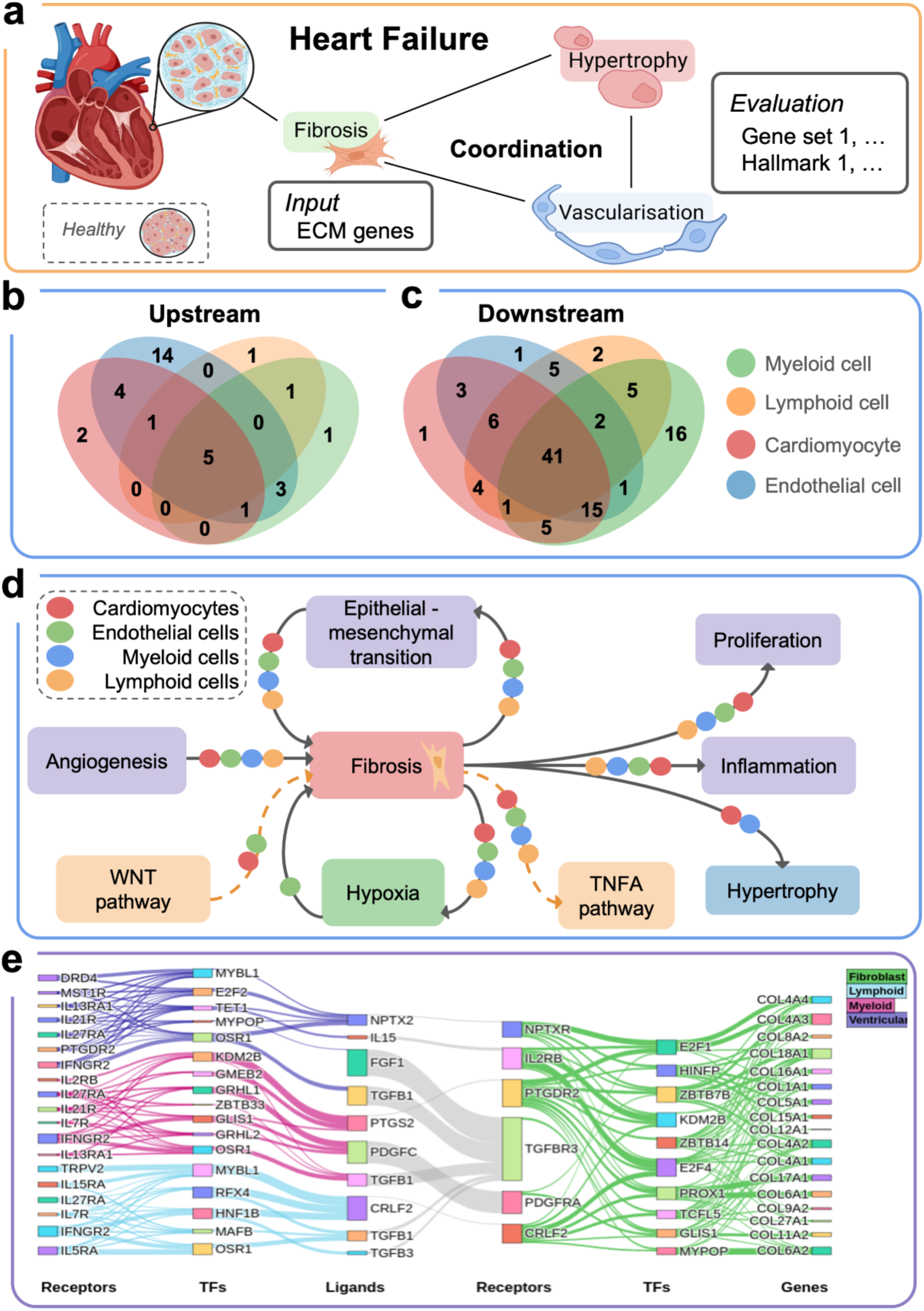
Analysis of intracellular regulations and multicellular coordination of cardiac fibrosis. **a)** Schematic representation of the cardiac fibrosis regulatory exploration with ReCoN. **b)** Venn diagram showing the intersection of gene sets associated with heart hypertrophy, vascularization, and fibrosis, found to be significantly enriched upstream of cardiac fibrosis in four major cell types: cardiomyocytes (red), myeloid cells (blue), lymphoid cells (orange), and endothelial cells (green). **c)** Venn diagram showing the intersections of gene sets significantly enriched downstream of cardiac fibrosis in the same four cell types. **d)** Schematic summary highlighting shared (purple) and cell type-specific (blue, green) biological programs identified as upstream or downstream of cardiac fibrosis. The symbols on the connecting arrows indicate the cell types involved. The most important identified pathways are additionally indicated in orange. **e)** Venn diagram showing the overlap between transcription factors (TFs) predicted by ReCoN (green) and those previously implicated in fibrosis (orange) or cardiac diseases (violet). Only the top 10 TFs were annotated from literature sources; full sizes of fibrosis- and cardiac disease-related receptor sets can therefore not be represented. **f)** Venn diagram showing the overlap between receptor sets predicted by ReCoN (green), a PKN network (blue), and receptors previously implicated in fibrosis (orange) or cardiac diseases (violet). As in panel e, only the top 25 receptors predicted by ReCoN or PKN have been annotated based on the literature. **g)** Sankey diagram showing a hierarchical organization of upstream regulators of the “NABA ECM collagens” gene set. Nodes are grouped by molecular type (e.g., transcription factors, receptors, ligands), and links represent the weighted, direct regulatory interactions present in the ReCoN-constructed HMLN. *Illustrations in a. were created using BioRender (*https://biorender.com).

Several known ligands mediators of cardiac fibrosis and remodeling, including TGFB1^42^, NPPB^43^, WNT5B^44^, BMP4^45^, GDF15^46^, and NRG1^47^, were retrieved as upstream regulators across multiple cell types (see Supplementary Notes 2). In contrast, we also predicted cell-type-specific genes related to cardiac fibrosis. In myeloid cells, the top-ranked gene was Oncostatin M (OSM), which was not prioritized in other lineages. Macrophages secrete OSM, which has been implicated in anti-fibrotic regulation^48^, supporting its potential role as a cell-type-specific paracrine signal during cardiac remodeling. In cardiomyocytes and endothelial cells, ADCYAP1 and FGF2 were specifically highly ranked. ADCYAP1 encodes PACAP, a neuropeptide involved in vasodilation with cardioprotective properties in HF^49–51^, while FGF2 is a well-known mediator of angiogenesis and cardiac repair^52^.

ReCoN also recovered several TF and receptors in fibroblasts that could regulate fibrosis activation from intracellular interactions (see Supplementary Notes 3, 4). Of note, ReCoN’s predictions suggest that MYPOP may be a key mediator of PFN-1’s effects in cardiac fibrosis and hypertrophy, aligning with its known molecular interactions. MYPOP, also known as *Myb-related transcription factor, Partner of Profilin-1* (PFN-1), interacts with PFN-1 and helps mediate its effects on gene expression^53^. While MYPOP has not been linked to heart failure yet, PFN-1 has been associated with cardiac hypertrophy and fibrosis^54^, and shown to contribute to fibrosis and cardiac injury in rat models^55^.

Overall, ReCoN effectively recovered many known regulators of fibrosis activation, both intracellular, with numerous TF and receptors, and ligands emitted by other cell types. Focusing on the ligand predictions, ReCoN identified regulatory signals broadly shared across cell types and cell-type-selective emitted signals regulating fibroblast activation and cardiac fibrosis.

### 2.6. Understanding multicellular programs in heart failure

Heart Failure (HF) emerges from a combination of cellular physiological alterations, including notably hypertrophy and vascularization remodelling. We used ReCoN to explore the relationship between these changes and fibrosis by predicting the sequence of events preceding and following fibroblast activation in HF.

ReCoN was used to retrieve both upstream regulators and downstream targets, which we integrated to draw a regulatory map of different HF hallmarks. From the predicted cell type profiles, we enriched several gene sets that illustrate these functions (see Supplementary Notes 7 and Supplementary Tables 8-11). These gene sets included general hallmarks and pathways related to hypertrophy and vascularization remodelling. We investigated the signals of cardiomyocytes, endothelial cells, lymphoid cells, and myeloid cells, as those four lineages showed the greatest improvement when including cell-cell communication and indirect effects in the Heart Failure showcase (see Figure 3g).

ReCoN identified several programs regulating fibroblast activation. Some were shared by the four cell types, including EMT, apical junction remodeling, and angiogenesis (see Figure 5b, Supplementary Table 8). These cellular processes underlie a general tissue remodelling process, preceding ECM protein production in cardiac fibrosis development (see Supplementary Notes 5). ReCoN also recovered cell-type-specific programs, particularly in cardiomyocytes and endothelial cells. Both cell types enriched the WNT signaling pathway, regulating the ECM gene expressions and fibroblast proliferation^42,56^. Endothelial cells uniquely strongly enrich hypoxia-related signaling, consistent with ischemia-driven vascular adaptation in HF (see Figure 5d, Supplementary Table 9).

In contrast, the post-fibrosis results enriched many gene sets across all lineages (41 out of 108 enriched gene sets), illustrating the global effect of cardiac fibrosis (see Figure 5c). These gene sets included several hallmarks of HF (see Figure 5d, Supplementary Table 10, Supplementary Notes 5), notably inflammation, hypertrophy, proliferation, hypoxia, EMT, and a set of genes specifically upregulated in systolic heart failure. Some interesting cell-type-specific gene sets were also enriched (Supplementary Table 11), notably both cardiomyocytes and myeloid cells enriched physiological and pathological hypertrophy pathways.

Together, these predictions from ReCoN decipher the coordination of several HF hallmarks and cellular processes with cardiac fibrosis. We summarized the specific and generic identified gene programs in Figure 5d. Angiogenesis seemed to precede fibrosis, while inflammation and hypertrophy were activated in response to fibrosis. Hypoxiad and EMT seemed to form positive feedback loop with fibrosis that may contribute to its maintenance and progression. Some of these pathways were shared across all cells, while others were cell-type-specific, highlighting the potential of ReCoN to identify both global and cell-type-specific mechanisms.

### 2.7. Visualisation of multicellular regulatory networks

Analysis with ReCoN relies on the exploration of the complete multicellular multilayer network. However, it can still be relevant to visualize complete receptor-TF-gene triplets to extract the most complete mechanistic hypotheses underlying ReCoN scores. We provide several functions to visualize the links between downstream genes and their regulators in each molecular layer. One can notably select the top TFs and receptors regulating a set of intracellular genes, and extract a subnetwork of direct connections between these elements. It is additionally possible to include the impact of other cell types through the top ligands predicted, and their upstream regulators, as presented in Figure 5e. In this example, we summarize the top regulators of the gene set “NABA ECM collagens”, used previously among genes involved in fibrosis.

## 3. Discussion

In this manuscript, we describe ReCoN, a method to reconstruct, analyze and visualize multicellular programs and cellular coordination to a molecular level.

ReCoN builds on previous approaches for multicellular programs by introducing mechanistic explanations of multicellular coordination, through multicellular modelling of microenvironments or tissues. The explicit network-based modelling of ReCoN provides interpretable and actionable insights across omics modalities, which combines both the intracellular details of classic GRN inference methods and the broad picture of CCC methods. The network exploration method in ReCoN allows predicting both regulators and consequences of perturbations across molecular layers. In any multicellular system, a signal can lead to the expression of secondary factors that reshape the coordinated behavior of neighboring cells. ReCoN is designed to understand the mechanisms of multicellular coordination, quantifying direct and environment-specific indirect effects of molecular signals. We envision ReCoN as a extension to prior multicellular modelling, offering an interesting compromise between prediction of cell type responses and understanding of their molecular coordination.

Our comprehensive evaluation demonstrated that each component of ReCoN, particularly the consideration of indirect effects, was useful in predicting the transcriptomic impact of individual or combinations of cytokines/ligands. ReCoN was used to model different microenvironments post-heart infarction and predicted distinct fibroblast trajectories in agreement with the literature. Finally, ReCoN proposed a comprehensive map of the multicellular coordination surrounding cardiac fibrosis, by recovering cell-type biological programs leading to/consequent to its activation.

ReCoN currently relies on transcriptomics-derived ligand–receptor information, which is known to be noisy, incomplete, and lacking spatial and protein-level constraints; such limitations likely cap the predictive gain we observe. Beyond ligand–receptor signaling, additional biological processes such as long-range paracrine effects, physical forces, extracellular matrix mechanics, or metabolic niches may not leave clear transcriptomic footprints and are thus underrepresented in the current. Moreover, evaluating ReCoN predictions remains challenging due to limited access to good-quality validation data. We argue with this work that the indirect effects might be an important component of molecular perturbation in multicellular contexts. Performance was indeed greatly improved when considering cellular coordination to predict cell transcriptomic effects. Additional data, notably intracellular perturbation in isolated and multicellular contexts, will enable more thorough validation of this hypothesis.

The modeling of multicellular systems still includes many challenges. ReCoN proposes a unified strategy for integrating molecular interactions across cell types and extracellular space. However, it can’t represent more than pairwise interactions or conditional ones. Borrowing conceptual representation from hypergraphs, which introduces edges involving more than two nodes, could be an extension to overcome this limitation. The network exploration implementation of ReCoN also present some limitations. While random walks with restarts offer a stable and fast exploration workflow for multilayer networks, it currently only considers positive weights to predict regulation strengths. It involves that the nature of the regulation, as activation or inhibition, has to be identified a posteriori.

In contrast to other modelling strategies, such as PhysiCell^57^ and agent-based frameworks^58,59^, our method does not simulate the dynamic evolution of tissues nor take into account individual cells’ spatial agencing. While these mechanistic models excel at capturing physical constraints, spatial organization, and precise cellular dynamics, they are currently less data-driven, scale less well to genome-wide molecular processes, and rely on hand-coded rules rather than learned regulatory dependencies. The development of specific network exploration methods, decomposing the graph exploration into smaller processes, could allow us to reconcile both dynamic and agent-based simulations and large multicellular multilayer networks.

## 4. Methods

### 4.1. Software and Reproducibility

The code of ReCoN is open-source and available at https://github.com/cantinilab/ReCoN. The notebooks and scripts used to generate the presented results are available at https://github.com/cantinilab/recon_reproducibility, along with the links to download the corresponding conda environments and singularity images.

### 4.2. Network Construction

ReCoN models context-specific regulation by integrating intracellular regulation with intercellular communication in a heterogeneous multi-layer network. A key requirement is that all edge weights remain strictly positive, ensuring valid probability distributions for diffusion algorithms. Only the nodes that are included in one of the layers are present in the final results, ignoring the ones present only in bipartites.

#### 4.2.1. Gene Regulatory Layer

We inferred a global gene regulatory network using HuMMuS (version 0.1.7), which combines scRNA-seq and scATAC-seq data to predict transcription factor–target gene interactions. However, any GRN inference method based on scRNA-seq alone or both scATAC-seq and scRNA-seq could be used.

With HuMMuS, we first built an HMLN composed of three layers: a TF layer, a scATAC layer containing peak co-accessibility information inferred from scATAC-seq data and the scRNA layer encoding transcriptional regulation inferred from scRNA-seq data. The co-accessibility network was inferred using CIRCE^60^ (version 0.3.4), and we kept all the links with positive scores. The scRNA layer was inferred with GRNBoost2^61^ (python version implemented in arboreto 0.1.6), and we kept the 50 000 first links. Both layers were then combined with the Python code of HuMMuS, and we explored the HMLN to compute the gene regulatory layer (see Supplementary Notes 6). Finally, only the links with a score above 1.5e-7 were retained in ReCoN’s gene regulatory layer.

#### 4.2.2. Receptor–Gene Bipartite Layer

While some took interest in inferring receptor-gene links directly^62^, there is very limited direct information compared to the large-scale ligand-target genes publicly available (Cytosig^63^, Nichenet^24^). We thus decided to infer receptor-gene links under the assumption that we reconstruct them through a linear matrix equation :

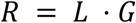

with ligand–receptor (*L*) and ligand–gene (*G*) adjacency matrices from NicheNet v2, accessible at https://zenodo.org/records/7074291. These matrices are accessible for both mouse (lr_network_mouse_21122021.rds, ligand_target_matrix_nsga2r_final_mouse.rds) and human (lr_network_human_21122021.rds, ligand_target_matrix_nsga2r_final.rds). We retrieved *R* using non-negative least squares (NNLS) in SciPy^64^ (v1.15.2), enforcing the production of only positive links as needed in ReCoN. We finally considered all receptor-gene links with a score above 0.005.

#### 4.2.3. Cell–Cell Communication Layer

The cell-cell communication layer contains nodes defined by a molecule and a cell type that produced it. Intercellular edges were inferred with the CellPhoneDB^65^ algorithm and the ligand-receptor database from Nichenet (https://zenodo.org/records/7074291) using LIANA+^8^ (version 0.1.9), without imposing a limit on the initial proportion of a cell type expressing a ligand (parameter expr_prop = 0). This produces a directed, weighted ligand–receptor network with each pair of cell types, using cell type scRNA-seq expression and a prior network of known ligand-receptor bindings. Any method producing non-negative edges can also be used as long as the links contain individual molecules and cell types involved. For the human data (Heart model), we used the “consensus” database of LIANA+ as prior. For the mouse data (Lymph node model), we use the ligand-receptor binding database provided by Nichenet for mice. We then retained all interactions strictly positive in all showcases.

### 4.3. Network exploration and signal propagation

#### 4.3.1. Random Walk with Restart (RWR)

Random walk with restart (RWR) is a stochastic process consisting of a succession of steps from one node (i.e., the seed) to a neighboring one through the network’s edges, with a probability to start again from the seed at each step. RWR can be used to explore HMLNs and to provide a measure of nodes’ closeness across the layers, ensuring the existence of a unique stationary distribution^25,66^. RWR strategies have been shown to significantly outperform methods based on local distance measures for the prioritization of gene-disease associations^18,67^. To run the RWR, we here used MultiXrank, a Python package proposing optimized RWR on universal multilayer networks^18^.

Different parameters guide the successive steps. First, the restart parameter determines a trade-off between returning to seed nodes and exploring the network and prevents the random walker from being trapped in dead ends. While the restart probability is typically 0.7 in previous works and HuMMuS^18,19,21,68^, the default value in ReCoN is 0.6 to allow a deeper exploration of the network across the large number of layers. If there are multiple seeds, each of them has a relative probability based on its weight to be used as a restart.

We also need to specify the probability of moving from one layer to any layer. It allows you to direct the flow of the exploration and the importance of each layer. By default in ReCoN, the probability of staying in a layer is 0.5. The probability of jumping to each reachable distinct layer is then 0.5/N, with N the number of reachable layers. Once the layer has been determined, the scores of the reachable nodes within it are normalized, allowing them to be used as probabilities for reaching them.

#### 4.3.2. ReCoN basic explorations

ReCoN has four standard ways of exploring multicellular systems, based on depth and direction. It can first be used to identify upstream and downstream molecules. The 2 exploration directions follow specific flows of information through different transition matrices in between layers. Downstream explorations allow transition from the cell-cell communication layer to the receptor layer, the receptor layer to the gene regulatory layer, and the gene regulatory layer to the cell-cell communication layer. It represents the expected signal transduction. Upstream explorations allow the exact opposite transitions. Additional layers can be included with their own rules, and allow transition back and forth with other layers (i.e., metabolites to genes and genes to metabolites).

For each exploration direction, it is possible to decide between intercellular exploration (exactly as described above) and intracellular exploration alone. The intracellular exploration excludes cell-cell communication and thus some transitions between layers. In the intracellular downstream exploration, there is no transition from the gene regulatory layers to the CCC layer: genes cannot jump to expressed ligands in the cell-cell communication layer. In the intercellular upstream exploration, transitions from the CCC layer to the gene regulatory layer are not allowed: binding ligands cannot jump to the gene that expresses them. Isolating both exploration depths is useful to understand the specific effect of direct signal transduction and the indirect effects through cell-cell communication.

#### 4.3.3. ReCoN predictions

The global effect or context of a perturbation is computed by a combination of its direct and indirect effects. The direct effect corresponds to an intracellular exploration, while the indirect effect is obtained from several intercellular explorations. All three outputs are vectors containing all the nodes of the HMLN and the associated probability to reach them. These probabilities are interpreted as the weight of the regulations between them and the seeds.

A downside of RWR in a large multilayer network is that nodes of layers that are close to the seeds have mechanically higher weights. Since the indirect effects modelling requires crossing more layers, the direct effect naturally has a much greater contribution. This is also a reason for differentiating direct and indirect effects, allowing for modulating in a second time their contribution. We additionally aim to reduce this effect in the computation of the indirect effect itself. From the direct effect, we first identify downstream genes in the GRN layer that encode secreted ligands. Rather than seeding the indirect RWR on those gene nodes, we assume that they will be translated as protein and seed the walk on the corresponding ligand-produced nodes in the cell–cell communication layer. This places the restart point closer to the receptors and downstream targets in other cells, reducing the score’s dilution across the walks. Symmetrically, for the upstream exploration, instead of starting from the binding ligands, we use the gene node linked to them.

#### 4.3.4. Cell type proportional contribution

Some cell types have more effect on their surroundings than others. It depends both on their distribution and their ligand expression profiles. By default, each cell type contributes equally in predicting the others. However, it is possible to adjust the Beta coefficient to represent it based on the available information for each dataset. Notably, spatial transcriptomic data could be used to identify the proximity and relative importance of each cell type over the sum of all surrounding cell signals.

### 4.4. Modeling Cytokine Treatments In Vivo (Mouse)

#### 4.4.1. Data Sources and Preprocessing

The Immune Dictionary data were downloaded at https://www.immune-dictionary.org as Seurat objects, and merged into a MuData object comprising the scRNA-seq profiles of 81 cytokine treatments across 17 lymph node cell types. In these downloadable files, a maximum of 100 cells per cytokine treatment for each cell type were sampled to ensure comparability across cell types for this analysis and 15 cell types were available: B cell, cDC1, cDC2, eTAC, ILC, Macrophage, Migratory DC, Monocyte, Neutrophil, NK cell, pDC, T cell (CD4+), T cell (CD8+), γδ T cell, Treg. All the cells were kept for the subsequent analysis. Only 41 cytokines were present in the prior ligand-receptor database, and 25 had at least one active connection inferred by LIANA+ with CellPhoneDB algorithm. We use the latter to compare the different models.

#### 4.4.2. External Multi-Omic Integration

To refine the gene regulatory layers, we combined scRNA-seq from external murine lymph nodes^69^ and scATAC-seq from murine immune cells (CD45+)^70^. The scATAC count matrix is accessible in GEO under accession code GSE242466 as “GSE242466_archr_mouse_immune_cell_atlas.tar.gz”. The scRNA-seq data are available at https://cf.10xgenomics.com/samples/cell-exp/7.2.0/4plex_mouse_LymphNode_Spleen_TotalSeqC_multiplex_LymphNode1_BC1_A_B1/4plex_mouse_LymphNode_Spleen_TotalSeqC_multiplex_LymphNode1_BC1_AB1_count_sample_filtered_feature_bc_matrix.h5, as part of the following dataset: https://www.10xgenomics.com/datasets/Mixture-of-cells-from-mouse-lymph-nodes-and-spleen-stained-with-totalseqc-mouse-universal-cocktail

Briefly, we filtered out in both matrices the features that were expressed in less than 3 cells, and then limited to only the 16,000 most variable genes. We then filtered out cells with fewer than 3 expressed features. It resulted in the scRNA-seq in 1,789 cells with 13,167 genes, and for the scATAC-seq in 3,759 cells with 254,545 regions.

#### 4.4.3. Murine lymph node model reconstruction

A gene layer was inferred from the external scRNA-seq datasets with GRNBoost2^61^ (python implementation in arboreto, v0.1.6), and a DNA region layer was inferred from the scATAC-seq dataset with CIRCE (v0.3.4). Untransformed counts were used as input for both. Both were then combined to infer a GRN with HuMMuS^21^ (v0.1.7). The cell-cell communication layer was computed from the Immune Dictionary subset of the cells treated with phosphate-buffered saline solution (PBS).

#### 4.4.4. Identification of Perturbed Genes

For each cytokine–cell type combination, differentially expressed genes were identified with Scanpy’s Wilcoxon rank-sum test (FDR P-val < 0.1, |log₂FC| > 1). In individual cytokine-cell type pairs evaluations, the ones with fewer than two significantly perturbed genes were excluded from downstream ranking analyses. We ended up with 206 cytokine-cell type pair profiles (see Supplementary Figure 2).

#### 4.4.5. Benchmarking Against Baseline Models

ReCoN models were compared to the ligand-PKN and receptor-PKN models, derived from the Nichenet database, on their ranking of the genes regulated by each cytokine. For the ligand-PKN model, it simply corresponds to the score of the ligand-gene matrix. For the receptor-PKN model, we multiplied the ligand-receptor and the receptor-gene matrix, equivalent to summing the regulatory score of the receptors bound by a cytokine, since the ligand-receptor matrix is unweighted.

For each model and cytokine, AUROCs were calculated comparing scores for individual cell types, then across cell types, to the list of perturbed genes established above. The multicellular ranking explores how individual cell type rankings are correctly scaled and comparable/mixed. It is an important criterion to retrieve cell-type-specific effects. However, it does not substitute for individual cell type rankings, because it can also mask cell type biases (see Supplementary Figures 5). For example, if a cell type A has twice as many differentially expressed genes as a cell type B, a model predicting all the genes of A above the genes of B could perform correctly, even if the genes in each cell type are poorly ordered. Studying the individual ranking can then unravel this bias by showing the gene order for each cell type. Statistical significance of performance differences was evaluated using two-sided Mann–Whitney U tests (non-Gaussian assumption), corroborated by Wilcoxon signed-rank tests.

### 4.5. Modeling Heart Failure in Humans

#### 4.5.1. Multiome Human Heart Cell Atlas Preprocessing

We preprocessed the single-cell multiome data from the human left ventricles as proposed in the GRETA pipeline. The samples were first extracted from the complete dataset. We filtered out the cells expressing fewer than 100 genes, and all genes expressed in fewer than 3 cells. We kept the most variable genes expressed in more than 16,384 cells, and the 65,536 most variable regions. Cell type annotations by different methods are already present in these samples. We kept all the cells whose annotations through unsupervised clustering, followed by marker gene annotations, through scANVI were coherent. The preprocessed dataset contained 25,787 cells, 16,384 genes, and 62,769 peaks.

#### 4.5.2. Human Heart model reconstruction

The gene regulatory layer, shared between all cell types, was then computed similarly to the murine lymph node model, using GRNboost (python v0.1.6) and CIRCE (v0.3.4), and inferring the final GRN combining both layers with the R version of hummus. Cell type identified from ReHeat2 was used. We thus matched cell types of the Heart Cell Atlas dataset to those of ReHeat2 according to the Supplementary Table 12. The cell-cell communication was then computed between these re-annotated cell types.

#### 4.5.3. Heart failure genes and ligands

To identify the cell-type-specific genes associated with HF, we used the MOFAcell scores of the multicellular factor 1 (MCP1) reported in ReHeat2^36^. To have high confidence sets of genes perturbed and unperturbed, we ranked all the scores and considered the top ten percent highest absolute scores as true positives, and the bottom ten percent as true negatives. In parallel, pairs of ligands and receptors with both associated with scores above an absolute gene loading of 0.1 were considered potential driver interactions in HF. The ligands of these pairs were used as seeds for ReCoN and PKN-ligands, while the receptors were used in PKN-receptors.

#### 4.5.4. Fibrosis Gene Selection

The genes related to the extracellular matrix were obtained from mSigDb. We merged all the NABA gene sets in human that were not related to cancer: ‘basement_membranes’, ‘collagens’, ‘core_matrisome’, ‘ecm_affiliated’, ‘ecm_glycoproteins’, ‘ecm_regulators’, ‘matrisome’, ‘secreted_factors’, ‘matrisome_associated’, ‘proteoglycans’. 149 of these genes were present in the gene regulatory layer and used as seeds in the fibrosis analysis showcases.

#### 4.5.5. Evaluation of performances in predicting HF differentially expressed genes

Seeds comprised HF-program ligands with normalized scores. RWR influence scores for all genes were computed as described above. We benchmarked against NicheNet’s ligand- and receptor-PKN models using AUROC/AUPR in multicellular and per–cell-type contexts.

#### 4.5.6. Evaluate receptor ranking shifts between ReCoN and PKN-receptors

We identify the receptor whose activity profile differs most between the ReCoN framework and the PKN-receptor model by treating each receptor’s activity score as a one-dimensional embedding representation and applying the Gene Mapping Matrix (GMM) approach^71^. Specifically, for each condition (ReCoN or PKN), we form a vector *U* of receptor scores and compute a normalized distance matrix

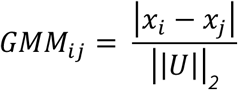

where ||*U*||_2_is the Euclidean norm of the score vector. We then quantify each receptor’s “movement” as the Euclidean distance between its corresponding row in the ReCoN GMM and its row in the PKN GMM. The receptor exhibiting the largest movement value is the one whose relative relationships to all other receptors shift most between the two models.

#### 4.5.7. Functional Enrichment

We used the gseapy Python package to realise the GeneSet Enrichment Analysis (GSEA) of upstream and downstream predictions. We evaluated the enrichment of the MSigDB Hallmarks (v2025) and the collection of gene sets extracted with the combination of keywords related to hypertrophy, vascularization, and fibrosis (see Supplementary Notes 7). Enrichment p-values were adjusted by Benjamini–Hochberg FDR. Adjusted P-values below 0.1 were considered significant in the subsequent analysis.

### 4.6. Modelling and comparing microenvironments of cardiac infarction

#### 4.6.1. Preprocessing of the spatial transcriptomics data

We deconvoluted the spatial data using the scRNA-seq reference^39^ and the cell2location package^72^. We kept the cells annotated as fibroblast, cardiomyocytes, endothelial cells, lymphoid cells, myeloid cells, vascular smooth muscle cells, or pericytes. With the *compute_expected_per_cell_type* function of cell2location, we then attributed a fraction of the spot gene counts to each cell type, which we then normalized on the total counts for each cell-type-spot profile. These cell-type-spot profiles were used later to create the specific cell-cell communication networks, and to calculate average for each spatial context the average cell type expression.

We then grouped the spots of the cell type niches 1, 4, and 5 annotations of the associated publication ^39^, associated with myogenic, fibrotic, and inflammatory contexts, respectively, on the slide Visium_18_CK296. This slide was chosen because it is located in the ischaemic regions and contains a high quantity of the three niches of interest. For each context, we grouped profiles of each cell type to obtain a context vector with gene expression * cell type as loadings, which will be used for defining a vector of seeds and their associated probabilities. We also used the cell type proportions per context calculated in the associated publication.

#### 4.6.2. ReCoN modelling and exploration

Each context impact was modeled with one RWR. The associated seeds and associated weights corresponded to the multiplication of the average gene expression per cell type, multiplied by the cell type proportion in each context. We only used the loadings of all cell types but the fibroblasts to consider the effect of the sole environment.

The cell-cell communication network was inferred using the cell-type-spot profiles and CellPhoneDB, in a similar way to the previous showcases, which used scRNA-seq data. We reused the GRN of the Heart Failure showcases, since both contexts contained the same cell types and the same receptor-to-gene links. We realised a downstream exploration with ReCoN to model the effect of the context on the fibroblast cells.

#### 4.6.3. Comparing responses through gene set enrichment

The profiles inferred by ReCoN were first very correlated in all three contexts. To focus on their differences, enrichments were done using the gseapy prerank function and the log+1 ratio of the context profile over the mean of the other context profiles. We used the standard MSigDB Hallmarks (v2025), such as in the cardiac fibrosis context, and lists of fibroblast state markers. For each fibroblast cluster identified in ^39^, markers were computed with t-tests (with Log Fold Change > 1 and p-value < 0.1 for thresholds) comparing the profiles of all cells belonging to one cell state with the rest of the cells of that cell type with scanpy’s rank_genes_groups function^73^ (v1.9.1).

#### 4.7. Data accessibility

Processed data and network used to generate the results will soon be available in Zotero.

Only public data was used to generate the results. The Immune Dictionary data can be downloaded at https://www.immune-dictionary.org/app/home. The murine lymph nodes scRNA-seq data and the murine scATAC-seq data of immune cells used as input for the lymph node model reconstruction can be downloaded on GEO under the accessibility number GSE242466 and on the 10Xgenomics platform https://www.10xgenomics.com/datasets/Mixture-of-cells-from-mouse-lymph-nodes-and-spleen-stained-with-totalseqc-mouse-universal-cocktail, respectively.

The Heart Human Cell Atlas data can be downloaded at https://cellgeni.cog.sanger.ac.uk/heartcellatlas/v2/Global_raw.h5ad (scRNA-seq) and https://cellgeni.cog.sanger.ac.uk/heartcellatlas/v2/Adult_Peaks.h5ad (scATAC-seq). We used the samples with the following IDs: HCAHeartST10773166_HCAHeartST10781063, HCAHeartST10773165_HCAHeartST10781062,H CAHeart9508627_HCAHeart9508819, HCAHeart9508629_HCAHeart9508821, HCAHeart9845431_HCAHeart9917173, HCAHeartST11064575_HCAHeartST11023240, HCAHeartST11064574_HCAHeartST11023239. The gene and ligand weights in the multicellular factor 1 identified in Heart Failure are available at https://doi.org/10.5281/zenodo.15261569. The gene sets were queried and downloaded from https://www.gsea-msigdb.org.

### 4.8. Code accessibility

ReCoN is publicly available at https://github.com/cantinilab/ReCoN. All the analyses presented in this manuscript were produced with ReCoN version 0.1.0 and will be uploaded to Zenodo. Notebooks and a conda environment to reproduce the analyses presented in this manuscript are available at https://github.com/cantinilab/ReCoN_paper.

Preprocessing and gene regulatory layer construction were made through Snakemake^74^ pipelines using Singularity^75^ containers. Downstream analyses are available as Jupyter Notebooks^76^. All computations were realised on the Pasteur Institute HPC, with 2 AMD EPYC 7552 48-Core Processors, 500 GB of RAM, and Linux Red Hat 8.8.

## 5. Author contributions

RT, LC, RORF, and JSR conceived the study. RT developed ReCoN and performed the computational experiments. RT and RORF searched and processed the data used in this study. LC and JSR supervised the project. RT, RORF, LC, and JSR wrote the manuscript. RT illustrated and organized the figures. All authors have read, edited, and approved the submission of the manuscript.

## 6. Acknowledgments

Computing was performed with the help of the HPC Core Facility of the Institut Pasteur. We thank Daniel Dimitrov for his advice regarding cell-cell communication and receptor signal transduction; and Jan Lanzer for his insights on multicellular communication, in particular in heart failure. The authors also thank Pau Badia i Mompel and the other developers of the GRETA pipeline for simplifying both the preprocessing of multiomics datasets and running gene regulatory network methods. We thank Zuzanna Słowik for her help in designing figures and proofreading the manuscript. We acknowledge the help of Deborah Philipps for administrative support.

