## Supplementary Notes, Figures for "Modelling multicellular coordination by bridging cell-cell communication and intracellular regulation through multilayer networks"

### Supplementary data

#### Supplementary Texts

##### Supplementary Notes 1. Cell type multilayer reconstruction.

Cell-type multilayers are individual heterogeneous multilayers, typically with each layer representing a different type of macromolecule. In the presented results, cell type multilayer was built from a GRN layer, which contains gene and TF, and a receptor layer. However, this framework is easily extendable. First, it is possible to add layers for new molecules. The only requirement is to provide at least one bipartite connecting the nodes of this new layer to the other layers. In the case of molecules with roles limited to intracellular regulation, the layer would be cell-specific and only connected to the layer of the cell-type they belong to. In the case where the layer provides intercellular bridges, it could also connect layers of different cell types. Second, it is possible to integrate layers between the macromolecules already represented in the default structure. For example, receptors could be linked both by ontology and by PPI relationships. In this case, the two layers could be considered as a multiplex. The RWR process can move freely and at no cost between the representations of the same node at each step, such as defined by transition probabilities specific to the multilayer. In other words, when exploring this layer, a fourth decision is added to the RWR process, where the layer that should be used for the next step is decided (i.e., the type of interactions).

##### Supplementary Notes 2. Cell-type-specific heart failure markers upstream of fibrosis extended results

TGFB1, NPPB, and WNT5B were consistently ranked in the top 50 out of 9,971 genes across all interacting cell types, suggesting broad relevance in fibrosis signaling. Other markers such as BMP4 and GDF15 showed selective prioritization, both ranked out of the top 1500 in lymphoid cells and in the top 20 for all other cell types, suggesting cell-type-specific involvement, even among canonical HF drivers.

**Supplementary Notes 3. ReCoN identified transcription factors related to heart remodelling and fibrosis**

ReCoN can also be applied to predict intracellular regulators, such as the TFs regulating the genes of the fibrosis hallmark. Among the top ten TFs predicted by ReCoN, six have been previously linked to fibrosis, and five to heart failure or related cardiac pathologies (see Supplementary Figure 4, Supplementary Table 3). MYPOP, FBXL10 (or KDM2B), and E2F1 were the three highest-ranked TFs. All three have been associated with both fibrosis and heart failure, suggesting they could play central roles in regulating cardiac fibroblast activation and ECM protein production. E2F1 has been proposed as a regulator of cardiac fibroblast differentiation[1] and has shown a cardioprotective role in right ventricular failure associated with pulmonary arterial hypertension[2]. FBXL10, an epigenetic regulator responsive to basic fibroblast growth factor[3], has been linked to several cardiac diseases[4], including diabetic cardiomyopathy[5]. This condition is often associated with interstitial fibrosis and can lead to heart failure. It has also been associated with fibroblast metabolic control[6] and appears to be upregulated during the intermediate stages of fibroblast-to-myofibroblast trans-differentiation[7].

MYPOP, also known as *Myb-related transcription factor, Partner of Profilin-1* (PFN-1), interacts with PFN-1 and helps mediate its effects on gene expression[8]. MYPOP has not been directly linked to heart failure. However, PFN-1 has been associated with cardiac hypertrophy and fibrosis[9], and shown to contribute to fibrosis and cardiac injury in rat models[10]. ReCoN's predictions suggest that MYPOP may be a key mediator of PFN-1's effects in cardiac fibrosis and hypertrophy, aligning with its known molecular interactions.

**Supplementary Notes 4. ReCoN leverages prior knowledge to predict receptors regulating ECM gene expression by cardiac fibroblasts.**

In addition to TFs, we used ReCoN to identify upstream receptors potentially driving ECM protein expression in cardiac fibroblasts. Predictions were compared to the receptor-PKN baseline, where each receptor was scored by summing its weighted links to target genes (see Supplementary Figure 4). Among the 25 top-scoring receptors prioritized by ReCoN, 23 were previously implicated in fibrotic processes across various tissues, and 16 could be specifically linked to cardiac fibrosis or related cardiovascular conditions (see Supplementary Table 4). ReCoN's top predictions showed strong agreement with the receptor-PKN, with 18 receptors shared between the two rankings. The prior knowledge model had comparable overlap with the literature, slightly higher with 24 receptors associated with fibrosis and 16 with cardiac-related contexts (see Supplementary Table 5).

Since the top receptors predicted by both methods had an important overlap, we next examined receptors whose scores differed the most between ReCoN and the receptor-PKN model, using a metric of "gene movement" across rankings[11]. While the novel receptors prioritized by ReCoN were not more specific to cardiac tissue, they remained enriched for regulators of fibrotic pathways (see Supplementary Table 6,7). Eight out of the top ten of these newly highlighted candidates were linked to fibrosis in the literature. This comparison also showed a higher ranking by ReCoN of several receptors previously described as cardiac-specific, refining their annotation to better reflect roles in fibrotic or remodeling processes.

These changes likely stem from ReCoN's integration of context-specific information, enabling it to highlight regulators relevant to the studied condition. However, this specificity may come at the cost of overlooking broadly validated receptors, making it difficult to assess whether the shifts represent true biological refinements or data-driven bias. In our analysis of receptor prioritization, ReCoN highlighted ADGRG1 (also known as GPR56) as a notable candidate, ranking it 12th compared to 50th in the receptor-PKN model. Recent studies have shown that cardiomyocyte-specific deletion of ADGRG1 leads to accelerated cardiac dysfunction, increased inflammation, and higher mortality, underscoring its potential role in heart failure pathogenesis. Conversely, IL1RAP, which ranked higher in the receptor-PKN model, was deprioritized by ReCoN. Blocking IL1RAP has been shown to reduce cardiac inflammation and preserve function in experimental models of myocarditis90.

Overall, ReCoN effectively recovered many known regulators of fibrosis, showing for the receptors a strong overlap with prior knowledge. At the same time, it reprioritized several receptors based on context-specific features of the data. These changes likely reflect a better adaptation to the studied condition but may also lead to the exclusion of broadly validated targets, making it difficult to assess whether the shifts represent true biological refinements or data-driven bias.

#### Supplementary Notes 5. Biological programs retrieved upstream and downstream of cardiac fibrosis extended results

Shared upstream programs included epithelial–mesenchymal transition (EMT), apical junction remodeling, and angiogenesis across all four cell types (Supplementary Table 8). EMT reflects the acquisition of mesenchymal traits and increased motility that underlie fibroblast activation and matrix deposition. On the other hand, apical junction remodeling indicates changes in cell–cell adhesion and polarity essential for tissue reorganization. Cardiac angiogenesis and fibrosis interact closely to promote cardiac regeneration[12].

Subsequently, the lineage-specific upstream enrichments emerged. Hypoxia-related signaling was strongly enriched only in endothelial cells, consistent with ischemia-driven vascular adaptation in HF[13] (see Figure 5d, Supplementary Table 9). Pathway gene sets for androgen response and progesterone inhibition were significant only in endothelial cells, albeit with lower NES, reflecting the specific role of sex-hormone modulation in vascular fibrosis. Cardiomyocytes uniquely enriched mTORC1 signaling, a key regulator of cell growth, protein synthesis, and metabolic adaptation under stress[14]. Additionally, both cardiomyocytes and endothelial cells enriched the mitochondrial pathways, reflecting high bioenergetic demand, and WNT signaling, which regulates extracellular matrix gene expression and fibroblast proliferation. All of which indicates a coordinated role in fibrotic remodeling.

Downstream of fibrosis, 41 out of 108 significant gene sets were shared across all lineages, despite showing different enrichment amplitudes. Shared axes corresponded to inflammation, hypertrophy, proliferation, hypoxia, and EMT (see Figure 5d, Supplementary Table 10), all important molecular hallmarks of HF. The inflammatory axis included TNFα/NF-κB, TGF-β, IFN-γ/α responses and generic inflammatory hallmarks, highlighting a global inflammation response and cytokine-driven remodeling. Hypertrophy programs, related to muscle hypertrophy and hypertrophic cardiomyopathy signatures, underline heart enlargement and contractile adaptation. Proliferation terms reflect cell-cycle activation in reparative and immune cells. Hypoxia and EMT remained significantly enriched downstream, marking their dual roles as drivers and consequences of fibrosis. Additionally, a gene set of upregulated genes in systolic heart failure was enriched across all lineages, with NES values varying from 1.97 (lymphoid) to 3.04 (cardiomyocytes). This range underlines a shared transcriptional response, while highlighting differences in activation strength among cell types.

**Supplementary Notes 6. GRN reconstruction with HuMMuS and hummuspy.**

HuMMuS is initially an R package, which relies on Python code for the RWR process, through the MultixRank package[15]. However, the computation of the individual layers can be computationally extensive in the default setting, since it relies on GENIE3 and Cicero. To work on bigger datasets, these layers can be computed externally (e.g., with Python packages), before computing the bipartites with the R version of HuMMuS. Once all the layers and bipartites are defined, a GRN can be obtained with both the R package or the Python code directly – *hummuspy*. The two versions provide the same results since they run the same Python code in the background. The R version was used for the Heart model, and the Python version was used for the Immune Dictionary. Since we started the development of ReCoN on the Immune Dictionary dataset, the Python version offered us easier integration with downstream analysis to define the default parameters of ReCoN.

**Supplementary Notes 7. A combination of keywords used to extract gene sets related to heart failure in MSigDB.**

The predictions of multicellular co-operation in heart failure and cardiac fibrosis were evaluated with gene set enrichments. We chose three categories of interest and extracted the related gene sets with the same keyword as in ReHeat2[16], which provide insights into multicellular co-operation in heart failure.

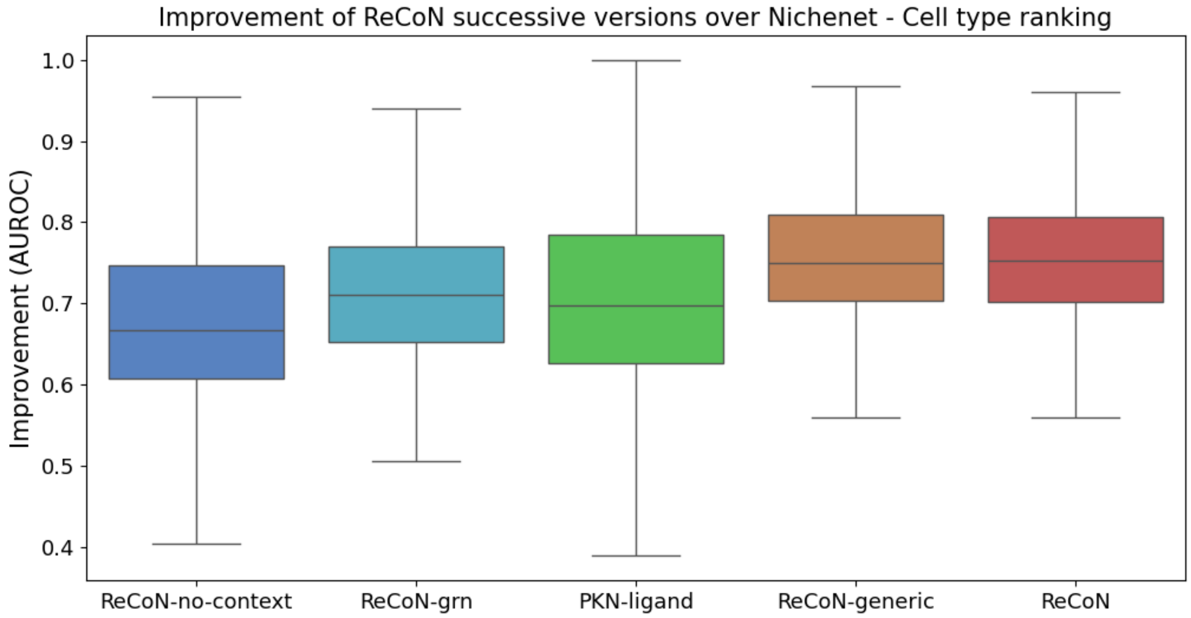


#### Supplementary Data

**Supplementary Figure 1. ReCoN predictions of transcriptomic responses across each cytokine and cell type pair – cell type ranking.** Four successively enriched versions of ReCoN (blue and green) and the PKN model from Nichenet are plotted. Boxplots represent the AUROCS of each model, where each cell type-cytokine pair considered is a value in the boxplot. Outliers are not plotted.

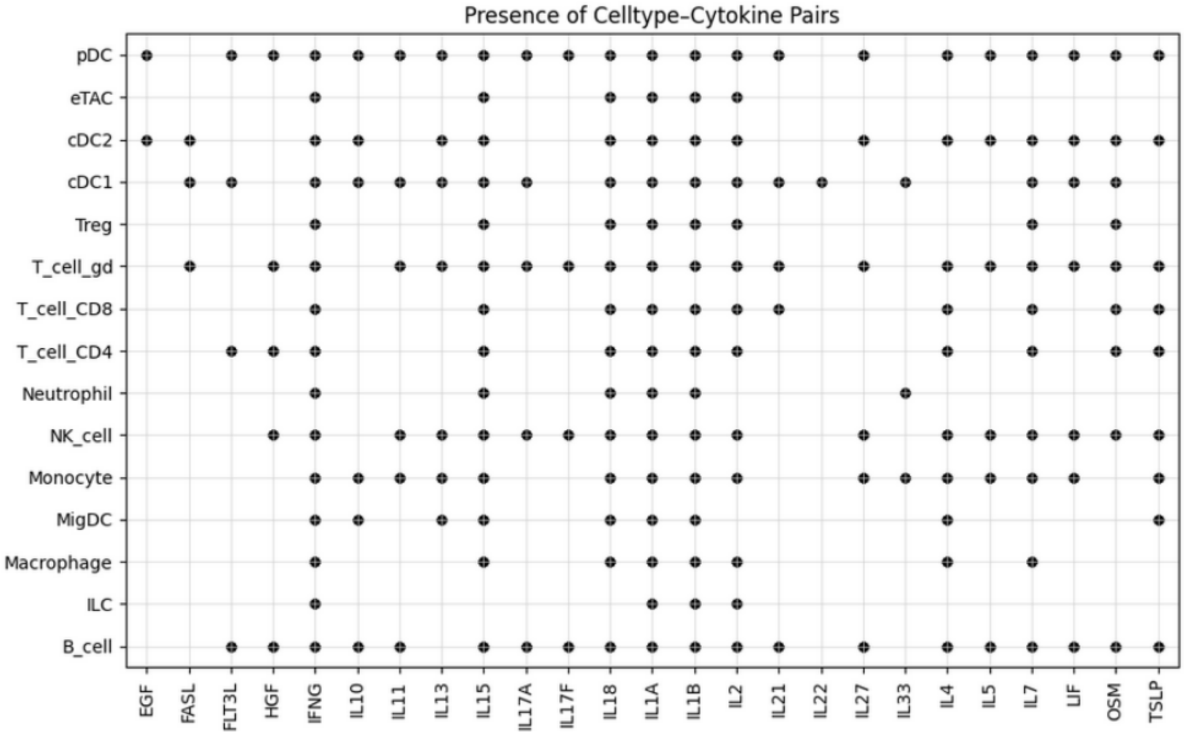

**Supplementary Figure 2. Pair of cytokines and cell types used in the study.** Only pairs with at least two significantly perturbed genes (see Methods) were considered here and for downstream analysis.

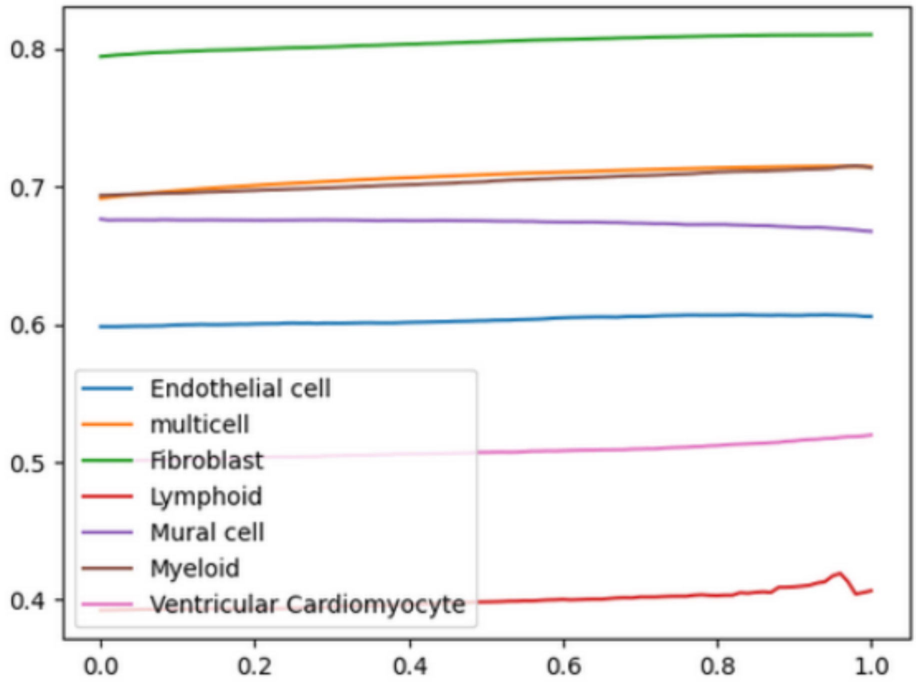


**Supplementary Figure 3. Cell type and multicellular AUPRC evolution depending on α.** AUPRC as a function of α (the weight of indirect effects) for individual cell type and global multicellular rankings in the HF showcase.

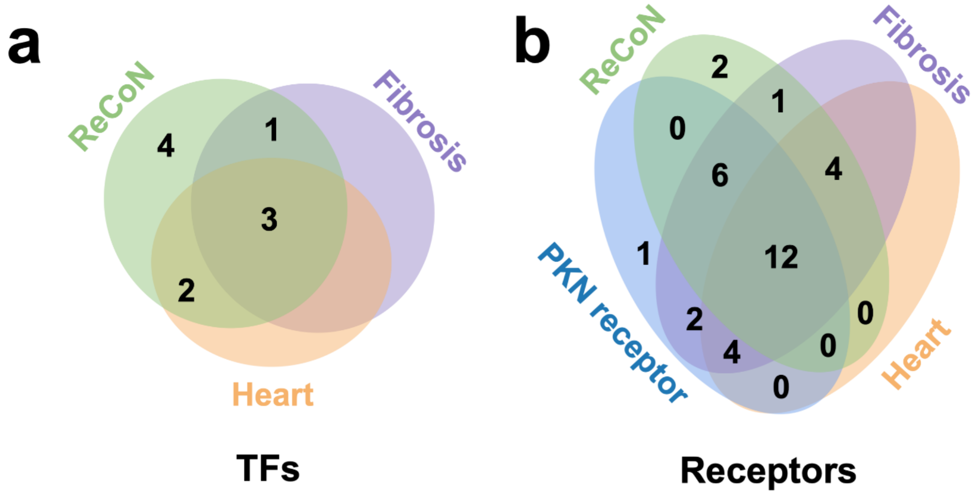


**Supplementary Figure 4. Overlap between predictions and literature for both TFs and receptors. a)** Venn diagram showing the overlap between transcription factors (TFs) predicted by ReCoN (green) and those previously implicated in fibrosis (orange) or cardiac diseases (violet). Only the top 10 TFs were annotated from literature sources; full sizes of fibrosis- and cardiac disease-related receptor sets can therefore not be represented. **b)** Venn diagram showing the overlap between receptor sets predicted by

ReCoN (green), a PKN network (blue), and receptors previously implicated in fibrosis (orange) or cardiac diseases (violet). As in panel a, only the top 25 receptors predicted by ReCoN or PKN have been annotated based on the literature.

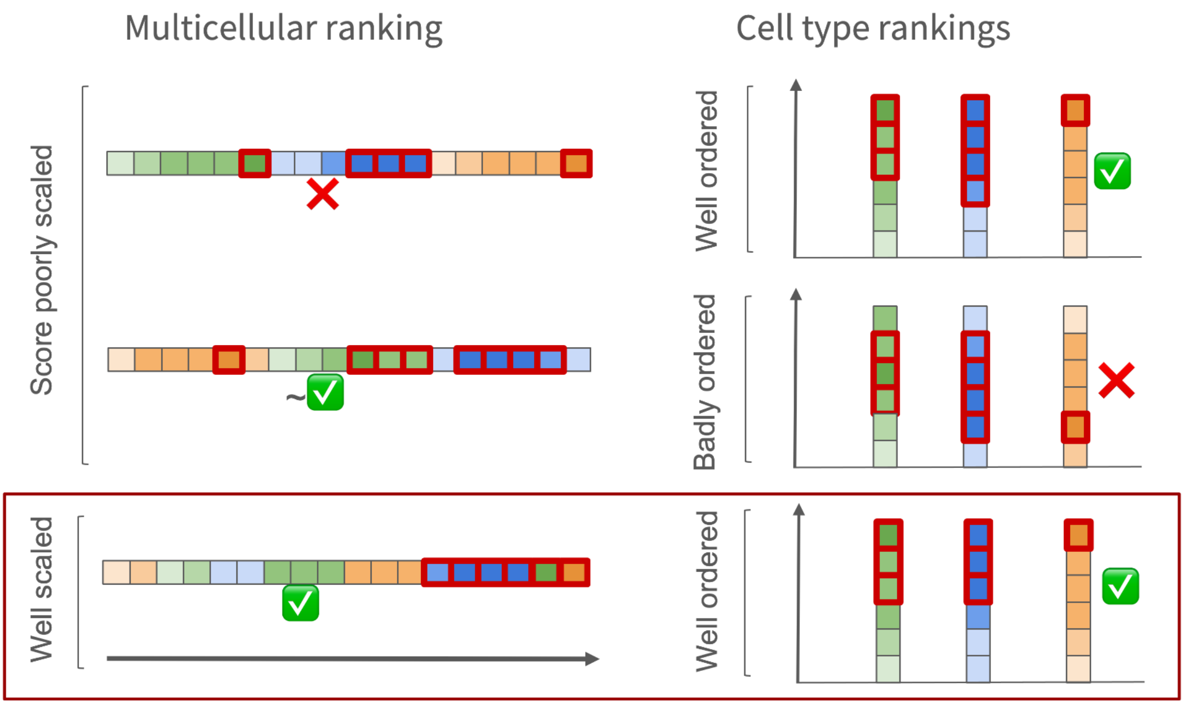


**Supplementary Figure 5. Complementarity between multicellular and cell-type gene rankings across different scenarios.** Each row illustrates a distinct scenario, showing the corresponding multicellular ranking (left) and cell-type–specific rankings (right). Squares represent genes ranked according to their predicted scores, with the color gradient indicating the true expression level. Red outlines highlight genes with the highest true expression across cell types. Scenario 1: Individual cell-type rankings are accurate with respect to true expression; however, their integration into a multicellular ranking is inaccurate. In particular, the cell type corresponding to the orange genes receives higher predicted scores, despite only one gene being truly highly expressed at the multicellular level. Scenario 2: The multicellular ranking appears accurate, as the blue cell type contains more highly expressed genes overall and these genes are correctly placed at the top of the multicellular ranking. However, cell-type rankings show poor discrimination of truly highly expressed genes within each cell type. Scenario 3: Both multicellular and cell-type rankings are accurate. The highest-expressed genes within each cell type are correctly ranked at the top of the corresponding cell-type rankings and are properly integrated into the multicellular ranking, accounting for differences in cell-type expression amplitudes. These examples illustrate that accurate multicellular inference requires both reliable cell-type–specific rankings and an appropriate integration across cell types

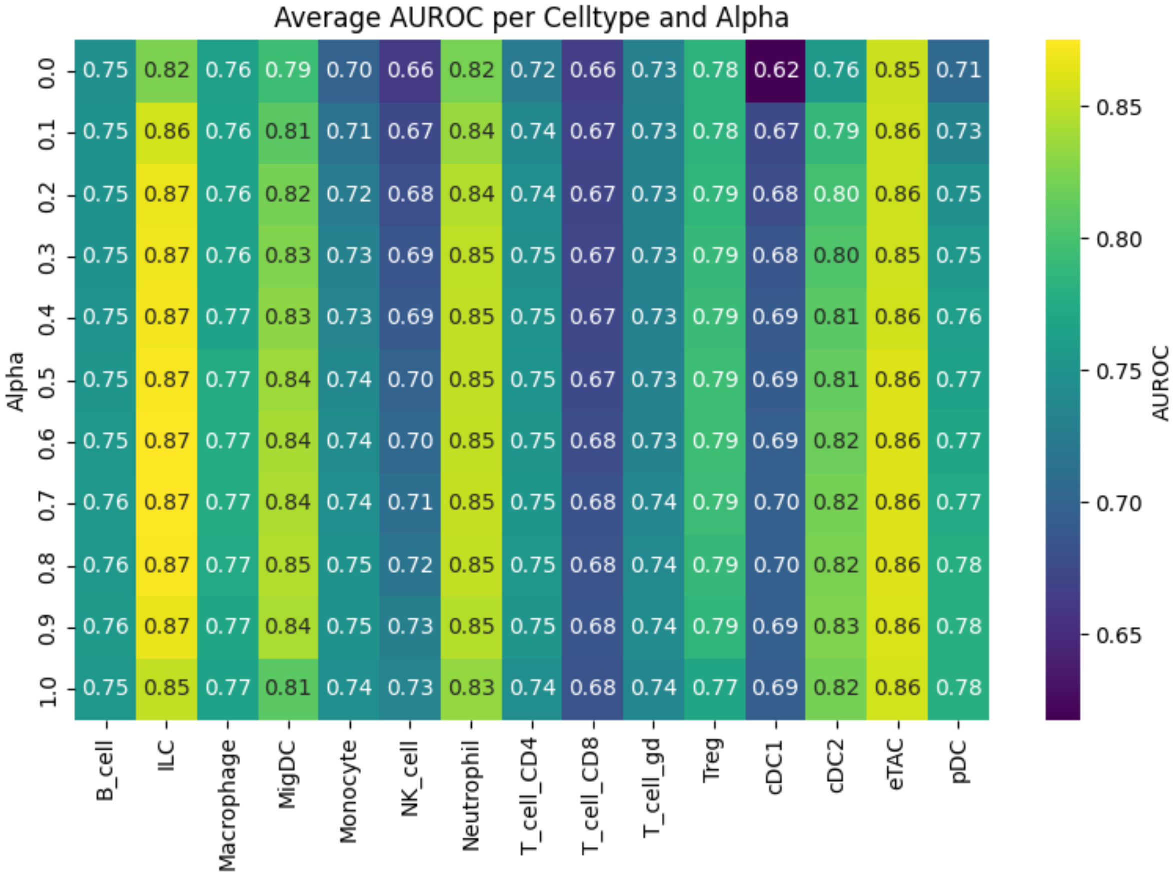


**Supplementary Table 1. AUROC across every cytokine and different α values – [immune dictionary showcase].**

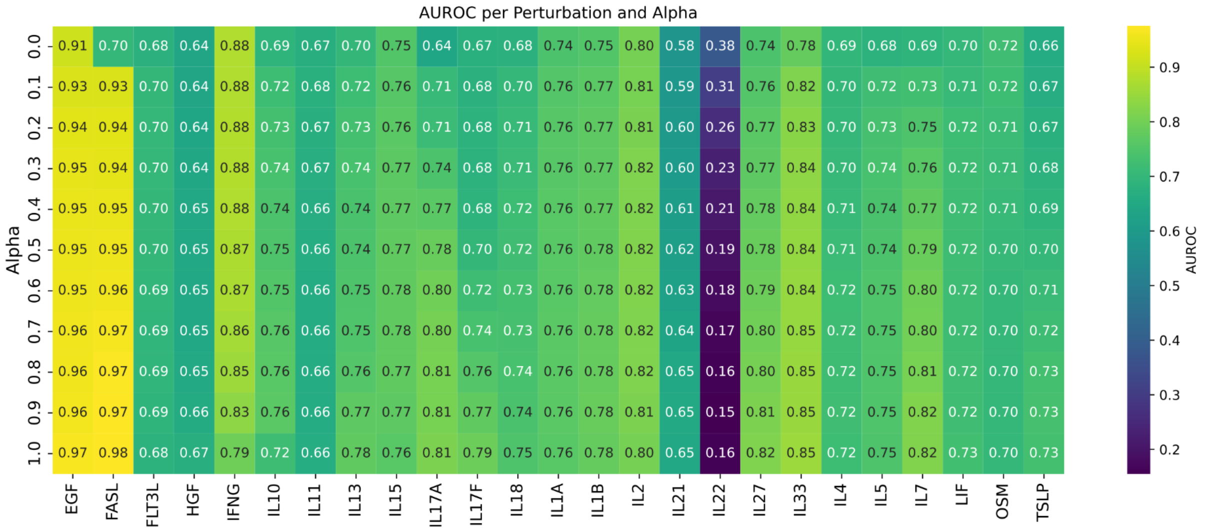


**Supplementary Table 2. AUROC across every cytokine and different α values – [immune dictionary showcase].**

| gene | score | fibrosis | heart | doi |
|---|---|---|---|---|
| **MYPOP** | 0.001065 | Related | Related | https://doi.org/10.1242/jcs.01618, https://doi.org/10.3892/mmr.2017.7446 |
| **KDM2B** | 0.001049 | YES | YES | https://doi.org/10.1111/jcmm.14146 |

| | | | | |
|---|---|---|---|---|
| **E2F1** | 0.000942 | YES | YES | https://doi.org/10.1161/circulationaha.120.047626, https://doi.org/10.1080/21655979.2021.1972194 |
| **PROX1** | 0.000921 | NO | YES | https://doi.org/10.1038/s41586-020-2998-x |
| **ZBTB7B** | 0.000915 | YES | NO | https://doi.org/10.1016/s0945-053x(01)00167-6 |
| **ZBTB14** | 0.000904 | YES | NO | https://doi.org/10.31083/j.fbl2809205 |
| **HINFP** | 0.000895 | NO | NO | — |
| **GLIS1** | 0.000834 | YES | YES | https://doi.org/10.1038/s41421-022-00490-3, https://doi.org/10.1016/j.freeradbiomed.2023.09.037 |
| **E2F4** | 0.000829 | YES | NO | https://doi.org/10.1096/fj.201903021rr |
| **TCFL5** | 0.000819 | NO | NO | — |

**Supplementary Table 3. Annotation of the 10 top TFs predicted by ReCoN – [heart failure showcase].** Receptors are classified as related to fibrosis and heart if a publication can justify this link. In violet are the receptors classified as related to both fibrosis and heart condition, in orange are the receptors related to one of the categories, and in red are the receptors related to neither of the categories.

| gene | score | fibrosis | heart | doi |
|---|---|---|---|---|
| **DRD4** | 0.005327 | YES | NO | https://doi.org/10.1111/acer.12047 |
| **IL27RA** | 0.004325 | YES | YES | https://doi.org/10.1016/j.heliyon.2023.e17099 |
| **IFNGR2** | 0.003781 | YES | YES | https://doi.org/10.1007/s10741-013-9393-8 |
| **PTGDR2** | 0.00351 | YES | YES | https://doi.org/10.1371/journal.ppat.1011812, https://doi.org/10.1016/j.yjmcc.2022.03.011 |
| **IL21R** | 0.00344 | YES | YES | https://doi.org/10.1016/j.ejphar.2022.175482 |
| **IL13RA1** | 0.003424 | YES | YES | https://doi.org/10.1161/JAHA.116.005108 |
| **GPR182** | 0.003365 | YES | YES | https://doi.org/10.1007/s10456-025-09977-5, https://doi.org/10.1161/jaha.117.007253 |
| **TRPV2** | 0.00332 | YES | YES | https://doi.org/10.1038/s41374-019-0349-z |
| **IL7R** | 0.003092 | YES | NO | https://doi.org/10.1172/JCI14685, https://doi.org/10.1186/s12931-022-02077-8 |
| **IL17RE** | 0.003079 | YES | Related | https://doi.org/10.3389/fcvm.2024.1470362 |
| **IL18RAP** | 0.002499 | YES | NO | https://doi.org/10.1002/hep.32776 |
| **ADGRG1** | 0.002461 | YES | YES | https://doi.org/10.1042/bsr20240826, https://doi.org/10.1177/1535370214529395 |
| **IL5RA** | 0.002353 | YES | Related | https://doi.org/10.1111/jcmm.18493 |
| **MST1R** | 0.002332 | YES | NO | https://doi.org/10.1111/liv.14892 |
| **IL15RA** | 0.002313 | YES | NO | https://doi.org/10.3389/fimmu.2024.1404891 |
| **CRLF2** | 0.00224 | YES | YES | https://doi.org/10.1096/fj.202302000rr |
| **PDGFRA** | 0.00221 | YES | Related | https://doi.org/10.1016/j.yexcr.2016.10.022 |
| **CD180** | 0.002201 | YES | YES | https://doi.org/10.1007/s00441-021-03488-7, https://doi.org/10.3892/mmr.2020.11242 |

| | | | | |
|---|---|---|---|---|
| **NPTXR** | 0.002105 | NO | NO | – |
| **TGFBR3** | 0.002091 | YES | YES | https://doi.org/10.1111/bph.13166 |
| **IL11RA** | 0.001949 | YES | YES | https://doi.org/10.1093/cvr/cvae224 |
| **MPL** | 0.001947 | NO | NO | – |
| **MUC5AC** | 0.001924 | YES | NO | https://doi.org/10.1371/journal.pone.0058658, https://doi.org/10.1038/mi.2012.114 |
| **IL12RB1** | 0.001906 | YES | YES | https://doi.org/10.3389/fphar.2020.00129 |
| **GPR25** | 0.001896 | YES | NO | https://doi.org/10.1111/febs.70117, https://doi.org/10.1016/j.jdermsci.2020.09.010 |

**Supplementary Table 4. Annotation of the 25 top receptors predicted by ReCoN – [heart failure showcase].** Receptors are classified as related to fibrosis and heart if a publication can justify this link. In violet are the receptors classified as related to both fibrosis and heart condition, in orange are the receptors related to one of the categories, and in red are the receptors related to neither of the categories.

| gene | score | fibrosis | heart | doi |
|---|---|---|---|---|
| **IFNGR2** | 1.849102 | YES | YES | https://doi.org/10.1007/s10741-013-9393-8 |
| **DRD4** | 1.749613 | YES | NO | https://doi.org/10.1111/acer.12047 |
| **IL13RA1** | 1.568187 | YES | YES | https://doi.org/10.1161/JAHA.116.005108 |
| **IL27RA** | 1.530149 | YES | YES | https://doi.org/10.1016/j.heliyon.2023.e17099 |
| **IL21R** | 1.240536 | YES | YES | https://doi.org/10.1016/j.ejphar.2022.175482 |
| **IL15RA** | 1.070769 | YES | NO | https://doi.org/10.3389/fimmu.2024.1404891 |
| **IL5RA** | 1.057473 | YES | Related | https://doi.org/10.1111/jcmm.18493 |
| **APCDD1** | 1.053718 | NO | NO | – |
| **TRPV2** | 1.031249 | YES | YES | https://doi.org/10.1038/s41374-019-0349-z |
| **PTGDR2** | 0.991871 | YES | YES | https://doi.org/10.1016/j.yjmcc.2022.03.011 |
| **IL18RAP** | 0.988052 | YES | NO | https://doi.org/10.1002/hep.32776 |
| **IL7R** | 0.982419 | YES | NO | https://doi.org/10.1172/JCI14685, https://doi.org/10.1186/s12931-022-02077-8 |
| **CD40LG** | 0.893819 | YES | YES | https://doi.org/10.1016/j.ijcard.2018.12.076 |
| **PDGFRA** | 0.852159 | YES | Related | https://doi.org/10.1016/j.yexcr.2016.10.022 |
| **MST1R** | 0.832997 | YES | NO | https://doi.org/10.1111/liv.14892 |
| **CRLF2** | 0.809771 | YES | YES | https://doi.org/10.1096/fj.202302000rr |
| **TGFBR3** | 0.767271 | YES | YES | https://doi.org/10.1111/bph.13166 |
| **IL1RAP** | 0.755057 | YES | YES | https://doi.org/10.1161/circheartfailure.124.011729 |
| **IL17RB** | 0.744823 | YES | NO | https://doi.org/10.1172/JCI14685, https://doi.org/10.1186/s12931-022-02077-8 |
| **IL10RA** | 0.675233 | YES | YES | https://doi.org/10.1161/CIRCULATIONAHA.117.027889 |
| **IL2RB** | 0.66985 | YES | YES | https://doi.org/10.1161/hypertensionaha.116.07084 |

| | | | | |
|---|---|---|---|---|
| **TNFRSF13C** | 0.649738 | YES | NO | https://doi.org/10.1126/sciadv.aas9944 |
| **GPR25** | 0.647856 | YES | NO | https://doi.org/10.1111/febs.70117, https://doi.org/10.1016/j.jdermsci.2020.09.010 |
| **CD180** | 0.624585 | Related | YES | https://doi.org/10.1007/s00441-021-03488-7 |
| **IL12RB1** | 0.591876 | YES | YES | https://doi.org/10.3389/fphar.2020.00129 |
| **MUC5AC** | 0.587984 | Related | NO | https://doi.org/10.1371/journal.pone.0058658 |

**Supplementary Table 5. Annotation of the 25 top receptors predicted by the PKN-receptor model – [heart failure showcase].** Receptors are classified as related to fibrosis and heart if a publication can justify this link. In violet are the receptors classified as related to both fibrosis and heart condition, in orange are the receptors related to one of the categories, and in red are the receptors related to neither of the categories.

| gene | score | ReCoN rank | PKN rank | fibrosis | heart | doi |
|---|---|---|---|---|---|---|
| **GPR182** | 37.167563 | 7 | 34 | YES | YES | https://doi.org/10.1016/j.jlr.2024.100679 |
| **IL17RE** | 31.16377 | 10 | 31 | YES | Related | https://doi.org/10.3389/fcvm.2024.1470362 |
| **ADGRG1** | 24.971621 | 12 | 50 | YES | YES | https://doi.org/10.1042/bsr20240826, https://doi.org/10.1177/1535370214529395 |
| **PTGDR2** | 17.204726 | 4 | 10 | YES | YES | https://doi.org/10.1371/journal.ppat.1011812, https://doi.org/10.1016/j.yjmcc.2022.03.011 |
| **NPTXR** | 12.26263 | 19 | 36 | NO | NO | – |
| **DRD4** | 12.252395 | 1 | 2 | YES | NO | https://doi.org/10.1111/acer.12047 |
| **CDH11** | 11.894496 | 50 | 105 | YES | YES | https://doi.org/10.3390/ijms24076549 |
| **LRP11** | 11.498974 | 48 | 94 | NO | NO | – |
| **TRPV2** | 11.183415 | 8 | 9 | YES | YES | https://doi.org/10.1038/s41374-019-0349-z |
| **CD180** | 10.82372 | 18 | 24 | YES | YES | https://doi.org/10.1038/cddis.2016.140, https://doi.org/10.1172/jci.insight.160684 |

**Supplementary Table 6. Annotation of the 10 top receptors with the highest gene movement ranked higher in ReCoN than in the PKN-receptor model – [heart failure showcase].** Receptors are classified as related to fibrosis and heart if a publication can justify this link. In violet are the receptors classified as related to both fibrosis and heart condition, in orange are the receptors related to one of the categories, and in red are the receptors related to neither of the categories.

| gene | score | ReCoN rank | PKN rank | fibrosis | heart | doi |
|---|---|---|---|---|---|---|
| **IFNGR2** | 25.24087 | 3 | 1 | YES | YES | https://doi.org/10.1007/s10741-013-9393-8 |
| **APCDD1** | 22.129364 | 28 | 8 | NO | NO | – |
| **IL13RA1** | 16.93084 | 6 | 3 | YES | YES | https://doi.org/10.1161/JAHA.116.005108 |
| **IL1RAP** | 16.331981 | 52 | 18 | YES | YES | https://doi.org/10.1161/circheartfailure.124.011729 |

| | | | | | | |
|---|---|---|---|---|---|---|
| **CD40LG** | 13.440412 | 29 | 13 | YES | YES | https://doi.org/10.1016/j.ijcard.2018.12.076 |
| **IL10RA** | 12.16169 | 54 | 20 | YES | YES | https://doi.org/10.1161/CIRCULATIONAHA.117.027889 |
| **OSMR** | 11.983381 | 68 | 35 | YES | YES | https://doi.org/10.1186/s12967-023-04163-x |
| **IL15RA** | 11.862556 | 15 | 6 | YES | NO | https://doi.org/10.3389/fimmu.2024.1404891 |
| **IL5RA** | 10.333617 | 13 | 7 | YES | Related | https://doi.org/10.1111/jcmm.18493 |
| **CAMK2A** | 7.037267 | 128 | 88 | YES | YES | https://doi.org/10.1016/j.jbc.2021.100893 |

**Supplementary Table 7. Annotation of the 10 top receptors with the highest gene movement ranked lower in ReCoN than in the PKN-receptor model – [heart failure showcase].** Receptors are classified as related to fibrosis and heart if a publication can justify this link. In violet are the receptors classified as related to both fibrosis and heart condition, in orange are the receptors related to one of the categories, and in red are the receptors related to neither of the categories.

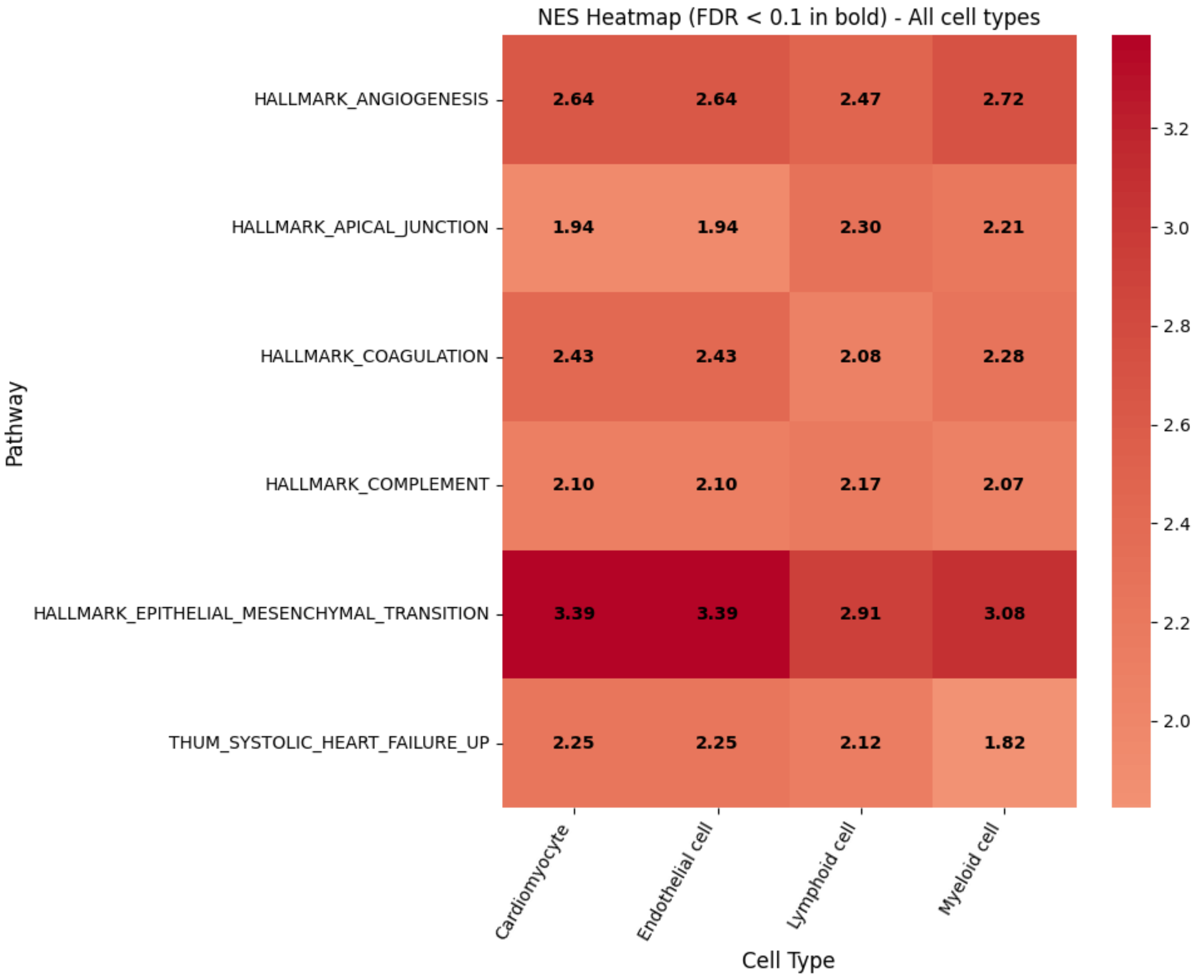


**Supplementary Table 8. Enriched pathways in all cardiac cell types upstream of fibrosis genes – [cardiac fibrosis showcase].** Here are all the gene sets enriched in all four cell types in the upstream exploration of section 2.6.

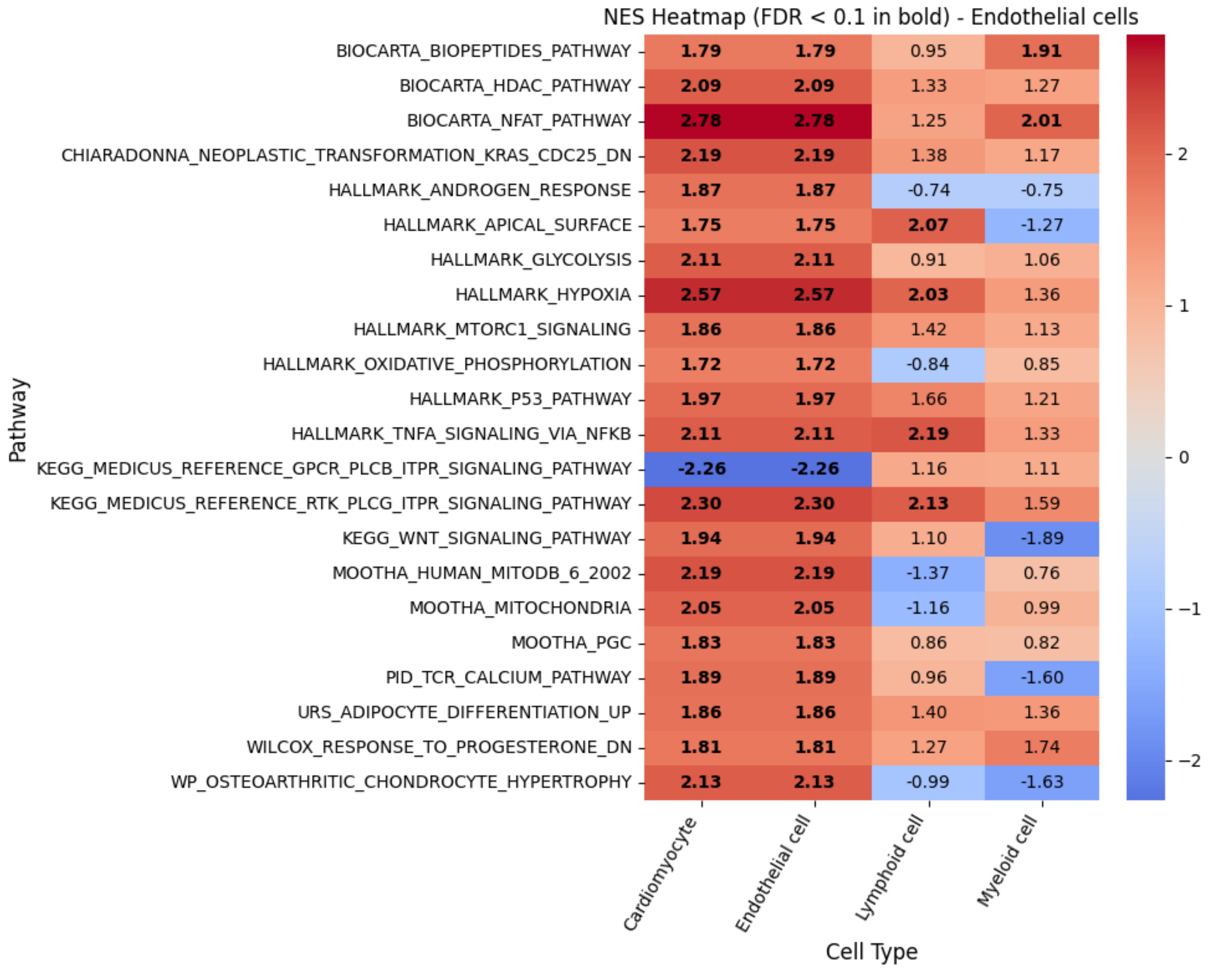


**Supplementary Table 9. Enriched pathways in endothelial cells upstream of fibrosis genes – [cardiac fibrosis showcase].** Here are all the gene sets enriched in Endothelial cells, in the upstream exploration of section 2.6.

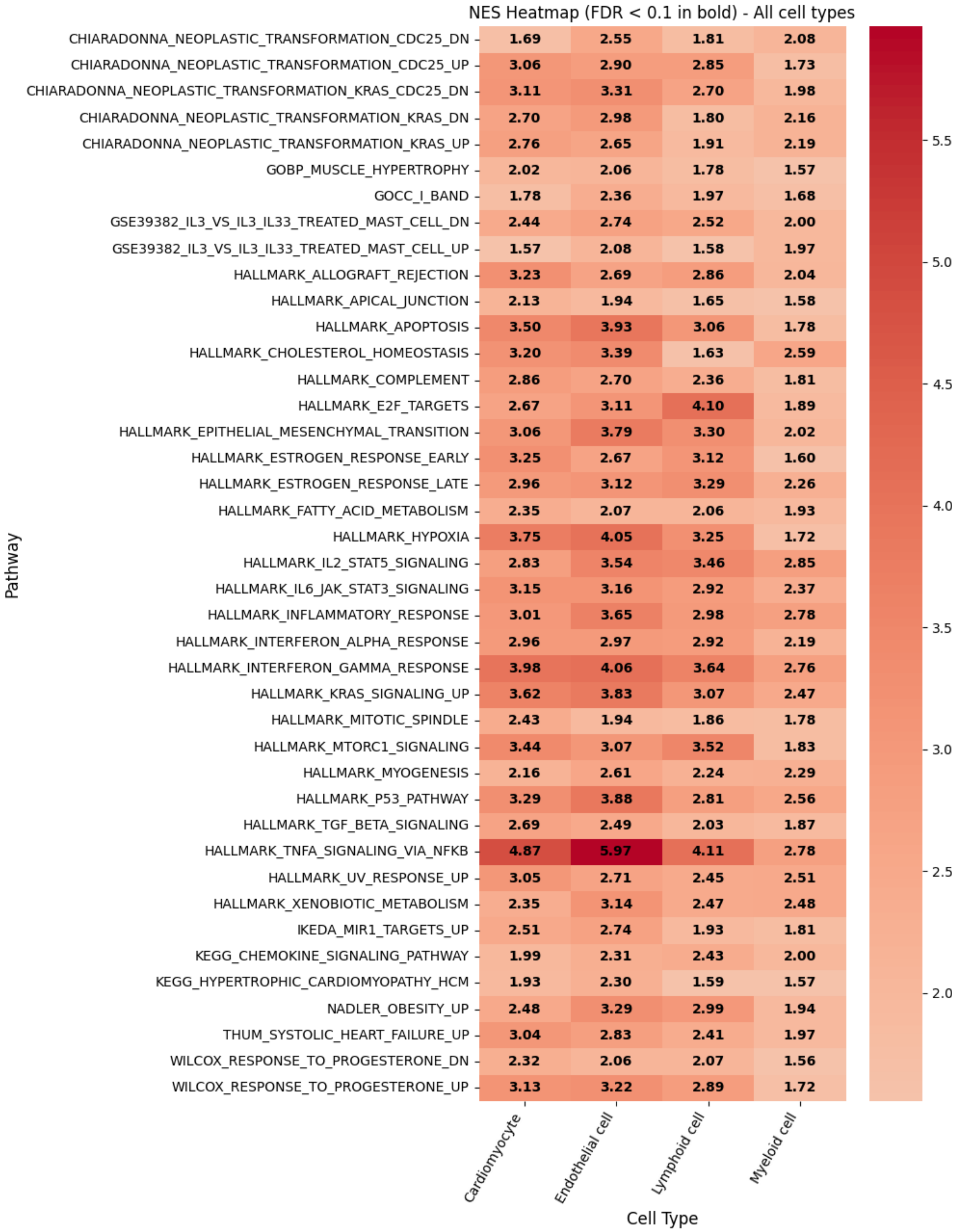


**Supplementary Table 10. Enriched pathways in all cardiac cell types downstream of fibrosis genes – [cardiac fibrosis showcase].** Here are all the gene sets enriched all four cell types considered in the downstream exploration of section 2.6.

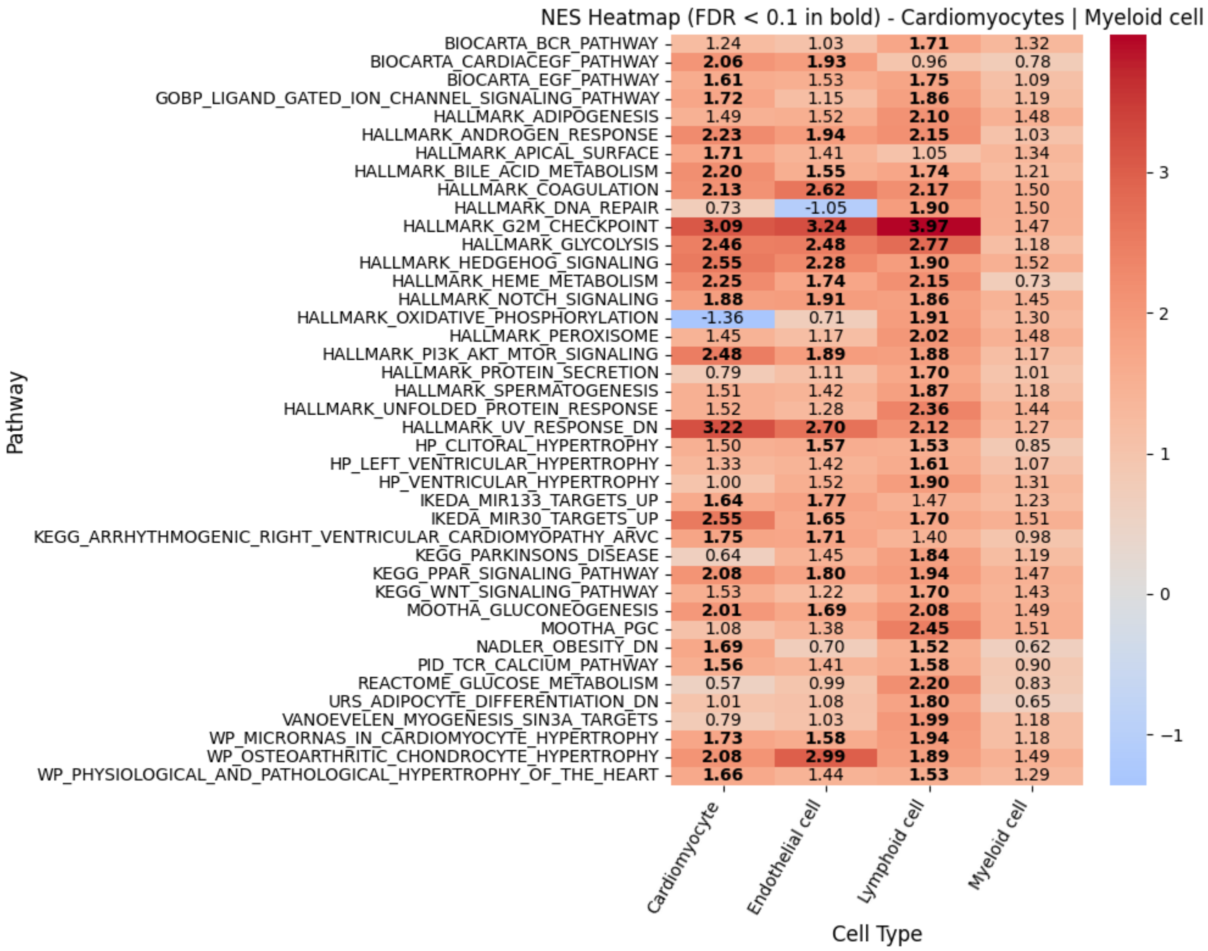

**Supplementary Table 11. Enriched pathways in cardiomyocytes and myeloid cells downstream of fibrosis genes – [cardiac fibrosis showcase].** Here are all the gene sets enriched in either cardiomyocytes or myeloid cells, and not in lymphoid cells, in the downstream exploration of section 2.6.

| Heart Cell Atlas | ReHeat2 - HF |
|---|---|
| CM | Ventricular Cardiomyocyte |
| Endo | Endothelial cell |
| Fib | Fibroblast |
| Lymphoid | Lymphoid |
| Myeloid | Myeloid |
| PC | Mural cell |
| vSMCs | Mural cell |

**Supplementary Table 12. Cell type matching between ReHeat2 and Heart Cell Atlas datasets – [cardiac fibrosis & heart failure showcase].** The left column corresponds to cell type annotations in the Heart Cell Atlas samples, the right column corresponds to the matching used in our model and in downstream analysis.
